# Fatal COVID-19 and non-COVID-19 Acute Respiratory Distress Syndrome is Associated with Incomplete Alveolar Type 1 Epithelial Cell Differentiation from the Transitional State Without Fibrosis

**DOI:** 10.1101/2021.01.12.426404

**Authors:** Christopher Ting, Mohit Aspal, Neil Vaishampayan, Steven K. Huang, Kent A. Riemondy, Fa Wang, Carol Farver, Rachel L. Zemans

## Abstract

ARDS due to COVID-19 and other etiologies results from injury to the alveolar epithelial cell (AEC) barrier resulting in noncardiogenic pulmonary edema, which causes acute respiratory failure; clinical recovery requires epithelial regeneration. During physiologic regeneration in mice, AEC2s proliferate, exit the cell cycle, and transiently assume a transitional state before differentiating into AEC1s; persistence of the transitional state is associated with pulmonary fibrosis in humans. It is unknown whether transitional cells emerge and differentiate into AEC1s without fibrosis in human ARDS and why transitional cells differentiate into AEC1s during physiologic regeneration but persist in fibrosis. We hypothesized that incomplete but ongoing AEC1 differentiation from transitional cells without fibrosis may underlie persistent barrier permeability and fatal acute respiratory failure in ARDS. Immunostaining of postmortem ARDS lungs revealed abundant transitional cells in organized monolayers on alveolar septa without fibrosis. They were typically cuboidal or partially spread, sometimes flat, and occasionally expressed AEC1 markers. Immunostaining and/or interrogation of scRNAseq datasets revealed that transitional cells in mouse models of physiologic regeneration, ARDS, and fibrosis express markers of cell cycle exit but only in fibrosis express a specific senescence marker. Thus, in severe, fatal early ARDS, AEC1 differentiation from transitional cells is incomplete, underlying persistent barrier permeability and respiratory failure, but ongoing without fibrosis; senescence of transitional cells may be associated with pulmonary fibrosis.

## Introduction

COVID-19 remains a global scourge, claiming thousands of lives daily and straining healthcare systems worldwide. In its most severe form, COVID-19 manifests as the acute respiratory distress syndrome (ARDS). COVID-19 ARDS causes acute hypoxemic respiratory failure with high mortality rates ^1–5^. However, the underlying mechanism by which COVID-19 ARDS results in refractory hypoxemia and high mortality is unknown.

Critical to the pathogenesis of ARDS is injury to the alveolar epithelium, which forms a tight barrier that maintains dry airspaces and permits gas exchange ^6^. The alveolar epithelium is comprised of AEC1s and AEC2s. AEC1s cover >95% of the alveolar surface, thus playing a critical role in barrier integrity, and are exquisitely thin, facilitating efficient gas exchange ^7, 8^. AEC2s are cuboidal, produce surfactant, and serve as progenitors ^9^. During ARDS, AEC death results in barrier permeability, leading to flooding of the airspaces with proteinaceous edema fluid, which in turn causes severe hypoxemia often necessitating mechanical ventilation, and death ^6^. Conversely, regeneration of the alveolar epithelium is critical for the restoration of barrier integrity, resolution of edema, liberation from the ventilator, and survival ^10, 11^. The state of epithelial injury and regeneration in COVID-19 ARDS has not been well characterized histologically. We speculated that incomplete alveolar regeneration may result in ongoing barrier permeability, pulmonary edema, and death from acute hypoxemic respiratory failure in COVID-19 ARDS.

The principal progenitor responsible for alveolar regeneration is the AEC2 ^9, 12–14^, though other progenitors may contribute ^15–17^. AEC2s proliferate and then spread and differentiate into AEC1s to restore normal alveolar architecture and barrier integrity. Using a mouse model of physiologic regeneration, we discovered that AEC2-to-AEC1 differentiation occurs in a nongradual manner ^18^. After proliferation, AEC2s exit the cell cycle and assume a transitional state before differentiating into AEC1s. The transitional state is characterized by markers of cell cycle exit, downregulation of AEC2 markers, a cuboidal or partially spread morphology, and expression of unique signature genes including Krt8. This state was subsequently identified in diverse mouse models of lung injury, suggesting that it is a conserved stage of regeneration regardless of the cause of injury ^19–23^. During physiologic regeneration in mice, the transitional state is transient; lineage tracing studies confirmed that transitional cells differentiate into AEC1s, restoring normal alveolar structure and function ^8, 18, 19, 24^. Which stage of alveolar regeneration - AEC2 proliferation, acquisition of the transitional state, or AEC1 differentiation - may be incomplete in COVID-19 is unknown. Autopsy studies of mechanically ventilated patients who died of acute respiratory failure have shown that the histology of COVID-19 ARDS is classic diffuse alveolar damage (DAD) and includes AEC2 hyperplasia ^25–29^. This suggests that proliferation may be preserved and that the prolonged barrier permeability and poor clinical outcomes in COVID-19 ARDS may be associated with incomplete conversion of AEC2s into the transitional state or differentiation of transitional cells into AEC1s.

Idiopathic pulmonary fibrosis (IPF) is caused by irreversible scarring of the lungs widely believed to arise from impaired alveolar regeneration after chronic or repetitive injury. We and others recently discovered that IPF is characterized by persistence of transitional cells with a paucity of mature AEC1s, suggesting that ineffectual differentiation of transitional cells into AEC1s may be the specific regenerative defect underlying the pathogenesis of fibrosis ^19–22, 24, 30, 31^. Since the transitional state has a transcriptomic signature of cell cycle arrest ^18, 19, 30, 31^, the persistence of transitional cells in IPF is consistent with emerging literature suggesting that epithelial cells in IPF are senescent ^32–39^. Recent case reports identified transitional AECs in the fibrosis that develops after months of nonresolving COVID-19 ARDS, so-called “fibroproliferative ARDS” ^40, 41^. In these cases, the transitional cells exist in a milieu of extensive matrix deposition and architectural distortion that was deemed irreversible, necessitating lung transplantation. In fact, the presence of transitional cells on biopsy was proposed as a potential biomarker of irreversible fibrosis ^40^. However, it is unknown whether transitional cells are inevitably associated with fibrosis in humans. Moreover, why transitional cells differentiate into AEC1s to restore normal alveolar architecture and function in mouse models of physiologic regeneration, whereas they persist and may beget fibrosis in human IPF and fibroproliferative ARDS, is a fundamental unanswered question in the field.

We hypothesized that in early (i.e., within the first 14 days of presentation) fatal COVID-19 ARDS, after severe epithelial injury, AEC2s proliferate and assume the transitional state, but that AEC1 differentiation from the transitional state is incomplete, resulting in ongoing barrier permeability, noncardiogenic pulmonary edema, ventilator dependence, and mortality. We further speculated that in early human ARDS, as in mouse models of physiologic regeneration, proliferating AEC2s exit the cell cycle and *transiently* adopt the transitional state but retain the capacity to differentiate into AEC1s, restoring normal alveolar architecture without fibrosis, whereas in IPF and fibroproliferative ARDS, after months of chronic or repetitive injury, AECs evolve into a state of *permanent* cell cycle arrest, or senescence, losing capacity for an AEC1 fate, and fibrosis ensues ^19, 30–35^. To explore these hypotheses, we obtained postmortem lung tissue of COVID-19 ARDS patients who died of acute respiratory failure within 14 days of presentation. For comparison, we obtained postmortem lung tissue from patients with early fatal ARDS of other etiologies and explanted lung tissue from patients with IPF undergoing lung transplantation. Tissue was examined for evidence of AEC2 proliferation, transitional cells, AEC1 differentiation, senescence, and fibrosis. Using existing single cell RNA sequencing (scRNAseq) datasets and immunostaining, we compared the gene expression profiles of transitional cells in two mouse models of physiologic regeneration without fibrosis, lipopolysaccharide (LPS) and pneumonectomy, early human COVID-19 ARDS, and human IPF, focusing on markers of cell cycle exit and senescence. The findings advance our basic understanding of physiologic and pathologic alveolar regeneration and have implications for clinical prognosis and management. Ultimately, investigation of the cellular and molecular mechanisms underlying ineffectual alveolar regeneration in ARDS and fibrosis may lead to novel therapies to promote physiologic regeneration, thus accelerating restoration of barrier integrity, resolution of edema, liberation from the ventilator, and survival in ARDS and preventing fibrosis in fibroproliferative ARDS and IPF.

## Materials and Methods

### Clinical Data and Tissue Acquisition

This study was approved by the University of Michigan Institutional Review Board. Patients were diagnosed with COVID- 19 by qPCR for SARS-CoV-2 on a nasopharyngeal, sputum, or BAL sample. Medical records were reviewed. Formalin-fixed, paraffin embedded lung tissue was obtained from the autopsies of COVID-19 patients who died at the University of Michigan Hospital. Normal lung from deceased individuals rejected for lung transplantation was obtained from Gift of Life Michigan. Idiopathic pulmonary fibrosis lung was explanted tissue from patients who underwent lung transplantation at the University of Michigan Hospital.

### Histology and Immunostaining

Sections were stained for hematoxylin and eosin (H&E) or trichrome and interpreted by a board-certified pathologist. For immunostaining, deparaffinized sections were boiled in Dako Target Retrieval Solution (Agilent S1699). The samples were blocked with 5% donkey serum in Tris-buffered saline with 0.05% Tween (TTBS), and incubated with antibodies against KRT8 (University of Iowa Developmental Studies Hybridoma Bank TROMA-1s, 1:20), HTI-56 (Terrace Biotech TB-29AHT1-56, 1:10), tomato lectin (Vector FL-1171, 1:250), α-smooth muscle actin-Cy3 (Sigma-Aldrich C6198, 1:200), pro-surfactant protein C (Millipore AB3786, 1:500), p53 (Abcam ab131442, 1:100), CDKN2A/p16INK4a (Abcam ab108349, 1:250), CCSP (Santa Cruz Biotechnology sc-365992, 1:100), and/or SARS nucleocapsid protein (Novus NB100-56576, 1:50-1:500) overnight at 4°C. Secondary antibodies, including FITC-conjugated anti-rat IgG (Jackson ImmunoResearch Laboratories 712-095-153, 1:250), Cy3-conjugated anti-rat IgG (Jackson ImmunoResearch Laboratories 711-167-003, 1:250), Cy5-conjugated anti-rat IgG (Jackson ImmunoResearch Laboratories 712-175-153, 1:250), Alexa-Fluor 555-conjugated anti-mouse IgG (Invitrogen A-31570, 1:250), Cy3-conjugated anti-rabbit IgG (Jackson ImmunoResearch Laboratories, 711-165-152, 1:250), and Cy5-conjugated anti-rabbit IgG (Jackson ImmunoResearch Laboratories, 711-175-152, 1:250) were applied for 1 hour at room temperature. For fluorescence *in situ* hybridization, sections were stained with SARS-CoV-2 probe (ACDBio #848561) as previously described ^18^. The slides were counterstained with DAPI 1:10000 in phosphate buffered saline and mounted on standard glass slides with Prolong Gold (Invitrogen P36930) antifade reagent. Images were acquired on an Olympus BX53 microscope or Aperio Leica scanner. Note that exposure settings were not always equal due to RBC autofluorescence but were set using isotype control antibodies. In all images, blue pseudocolor is DAPI.

### Analysis of scRNAseq Datasets

Single cell RNA sequencing datasets were interrogated for gene expression levels. Datasets were obtained from the NCBI Gene Expression Omnibus (https://www.nci.nlm.nih.gov/geo) under accession numbers <u>GSE113049</u> ^18^, <u>GSE138585</u> ^21^, <u>GSE136831</u> ^30^, <u>GSE106960</u> ^42^, and <u>GSE135893</u> ^31^. UMI count matrices were reprocessed to obtain average log normalized expression values for each cell type annotated in the respective study. In the mouse LPS scRNAseq study, mice had been treated with 45 μg LPS intratracheally and euthanized at 7 days later . In the pneumonectomy scRNAseq study, mice had been subjected to pneumonectomy of the right lobe and euthanized 7 or 21 days later ^21^. In both studies, lineage-labeled AEC2s were sequenced. In the human IPF studies, the lungs of IPF patients had been subjected to scRNAseq ^30, 31^. The AEC2 state was identified as “naïve AEC2s” from the LPS dataset ^18^, subpopulation II from the pneumonectomy dataset ^21^, and AEC2s from the human IPF datasets ^30, 31^. The transitional state was identified as the “cell cycle arrest” cluster in the LPS dataset ^18^, “subpopulation I” from the pneumonectomy day 7 wildtype mice ^21^, and the “aberrant basaloid” ^30^ and “KRT5-/KRT17+” ^31^ clusters from the human IPF datasets. The AEC1 state was identified as “naïve AEC1s” from the LPS dataset ^18^, AEC1s from a dataset used as a control for the pneumonectomy dataset ^42^, and AEC1s from the human IPF datasets ^30, 31^.

## Results

### Clinical Presentation

All ARDS patients presented with acute hypoxemic respiratory failure and radiographic evidence of pulmonary edema (Supplemental Figure 1, Supplemental Tables 1,2). All patients died within two weeks of hospitalization. Patients #1-3 were diagnosed with COVID-19 by SARS-CoV-2 PCR. Patients #4-6 were hospitalized in 2018. Clinical autopsy reports revealed DAD.

### Severe Epithelial Damage in COVID-19 ARDS

To characterize epithelial injury in COVID-19 ARDS, we first examined histologic lung sections stained with H&E. Consistent with prior reports ^25–28^, the histology of COVID-19 ARDS was acute DAD. There was diffuse airspace filling with edema, fibrin, and hyaline membranes (Figure 1A-D), indicative of increased epithelial permeability. Interstitial and alveolar inflammation, predominantly macrophages and lymphocytes, were present. Desquamated epithelial cells with elongated morphology were observed (Figure 1D). Such cells are commonly designated AEC1s ^43^ but are often, and in this case were, thicker and cover less surface area than AEC1s ^44^, raising new doubt as to their identity. Regardless, the radiographic and histologic findings observed in COVID-19 ARDS indicated epithelial injury. To specifically assess the extent of structural damage to AEC1s and AEC2s, we stained sections for AEC1 and AEC2 markers. AEC1 markers appeared diffuse and speckled throughout the lungs, consistent with extensive AEC1 injury (Figure 1E,F). The septa were thickened, likely due to the presence of edema and inflammatory cells in the interstitial space (asterisks in Figure 1E,F). Small defects in staining were frequent (Figure 1E). Complete absence of staining from one or both sides of the alveolar septa were present but infrequent (Figure 1F) and were interpreted as AEC1 loss resulting in denuded septa. Staining for the AEC2 marker pro-surfactant protein (SP) C revealed vast areas devoid of mature (proSPC+) AEC2s with only rare proSPC+ AECs remaining (Figure 1G, Supplemental Figure 2). The near complete absence of SPC+ cells may indicate AEC2 death or SPC downregulation. Taken together, the clinical, radiographic, and histologic data indicated severe, extensive epithelial damage, resulting in noncardiogenic pulmonary edema, in turn leading to acute hypoxemic respiratory failure.

**Figure 1.**
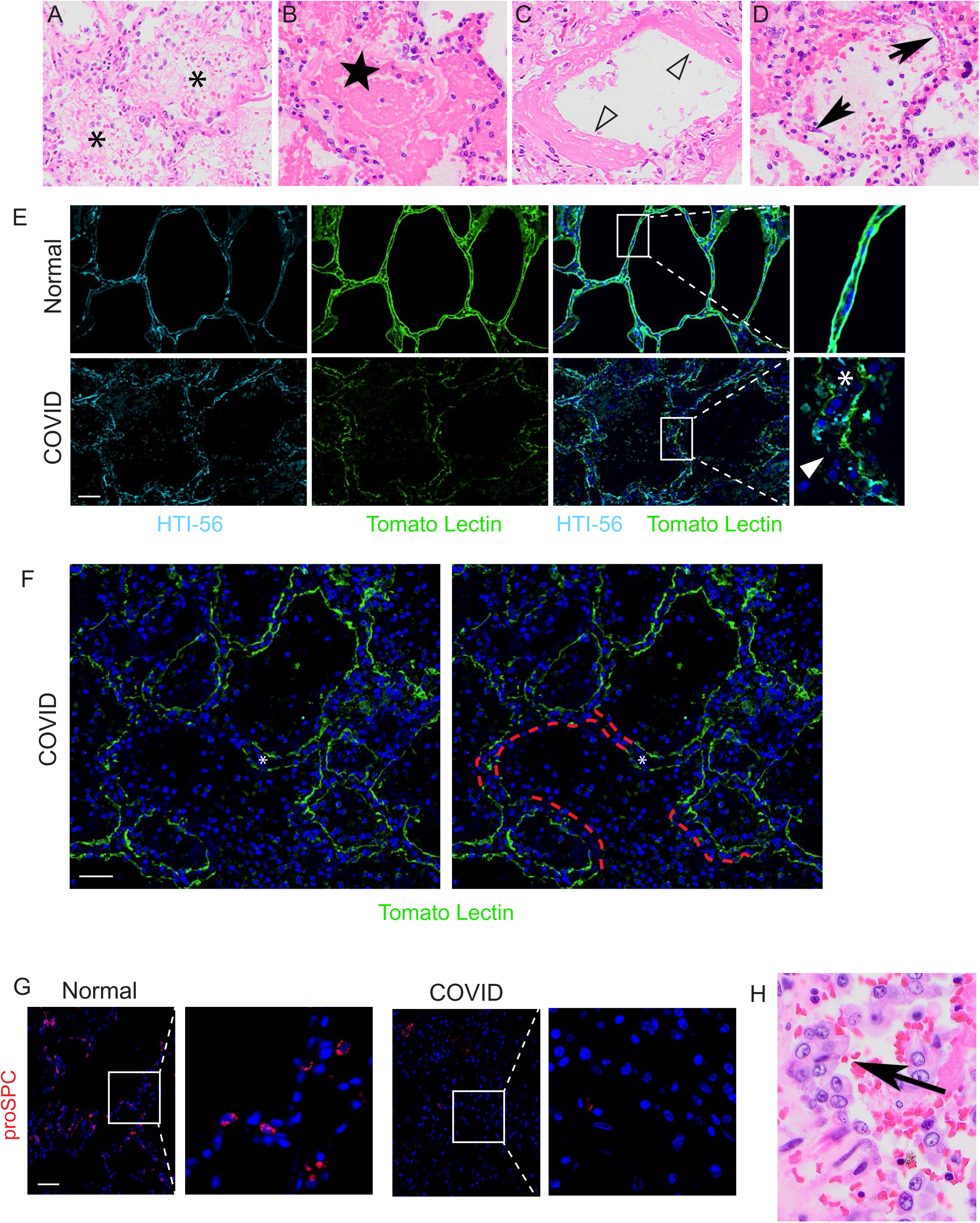
Epithelial Injury and Proliferation in COVID-19 ARDS. H&E staining of lungs from COVID-19 patients reveals (A-D) acute DAD with (A) edema (asterisks), (B) fibrin (star), (C) hyaline membranes (open arrowheads), and (D) desquamated epithelial cells (arrows) (200x). E) Immunostaining for the AEC1 markers HTI-56 and tomato lectin reveals a speckled pattern diffusely with frequent small defects in staining (arrowhead) and septal thickening consistent with interstitial edema and/or inflammation (asterisk). Scale bar = 50 µm. F) Immunostaining for the AEC1 markers HTI-56 and tomato lectin reveals occasional areas in which staining was completely absent from one or both sides of the alveolar septa (indicated by red dashed tracing), consistent with denudation. Septal thickening indicated by asterisk. Scale bar = 50 µm. G) Immunostaining for the AEC2 marker proSPC revealed vast areas devoid of mature AEC2s with rare AEC2s remaining. Scale bar = 50 µm. H) H&E revealed hyperplastic cuboidal epithelial cells (arrow, 40x). n=3.

### AEC2s Proliferate and Assume the KRT8^hi^ Transitional State in COVID-19 ARDS

To assess whether surviving AEC2s are mobilized to regenerate the injured epithelium, we again examined histologic sections stained with H&E. We observed hyperplastic cuboidal epithelial cells (Figure 1H), a common histologic feature of DAD which has historically been termed “AEC2 hyperplasia” ^43^. Indeed, SPC staining did reveal rare hyperplastic AEC2s (Supplemental Figure 3). The hyperplastic AEC2s were hypertrophic, consistent with previous observations in mouse models of lung injury ^45^. However, hyperplasia of mature (proSPC+) AEC2s was exceedingly rare amidst vast areas devoid of mature AEC2s (Figures 1G, Supplemental Figure 2). This suggests that the majority of the hyperplastic epithelial cells observed by H&E staining do not retain a mature AEC2 phenotype. Regardless, the presence of hyperplastic cuboidal epithelial cells on H&E suggested that AEC2s (or other progenitors) had successfully proliferated in attempt to replace damaged AECs.

To determine whether regenerating AEC2s were able to assume the transitional state, we stained sections for the transitional state marker KRT8. KRT8^hi^ transitional cells were abundant throughout the COVID-19 lungs but absent in normal lungs (Figure 2A,B, Supplemental Figure 4). [AEC1s and AEC2s in normal lungs express KRT8 at much lower levels than transitional cells in injured lungs (Supplemental Figure 5) ^18–22, 24, 30, 31^.] Some transitional cells were cuboidal; some existed as single cells while others were doublets or hyperplastic suggesting recent cell division (Figure 2A, Supplemental Figure 4). However, the morphology of transitional cells was often partially spread and occasionally flat approaching AEC1 morphology (Figure 2A, Supplemental Figure 4). The cuboidal hyperplastic, partially spread, and flat transitional cells typically existed as monolayers along structurally normal septa (Figure 2A, Supplemental Figure 4), consistent with organized proliferation and ongoing differentiation into AEC1s. Occasional transitional cells were hypertrophic, bizarrely shaped, and disorganized (Figure 2A, Supplemental Figure 4). KRT8^hi^ transitional cells did not express SPC (Figure 2B), consistent with known downregulation of AEC2 markers as cells assume the transitional state^18–22, 30, 31^. [Although transitional cells may arise from club-like progenitors ^15, 20, 30, 31^, bronchiolization was not observed, and transitional cells did not express the club cell marker CCSP (Supplemental Figure 6).]. Taken together, these data demonstrate that transitional cells arise within the first two weeks of onset of human ARDS. The abundance of transitional cells (Figure 2A, Supplemental Figure 4) suggests that the paucity of SPC+ cells may be attributable to both AEC2 death due to cytopathic viral infection and successful acquisition of the transitional state.

**Figure 2.**
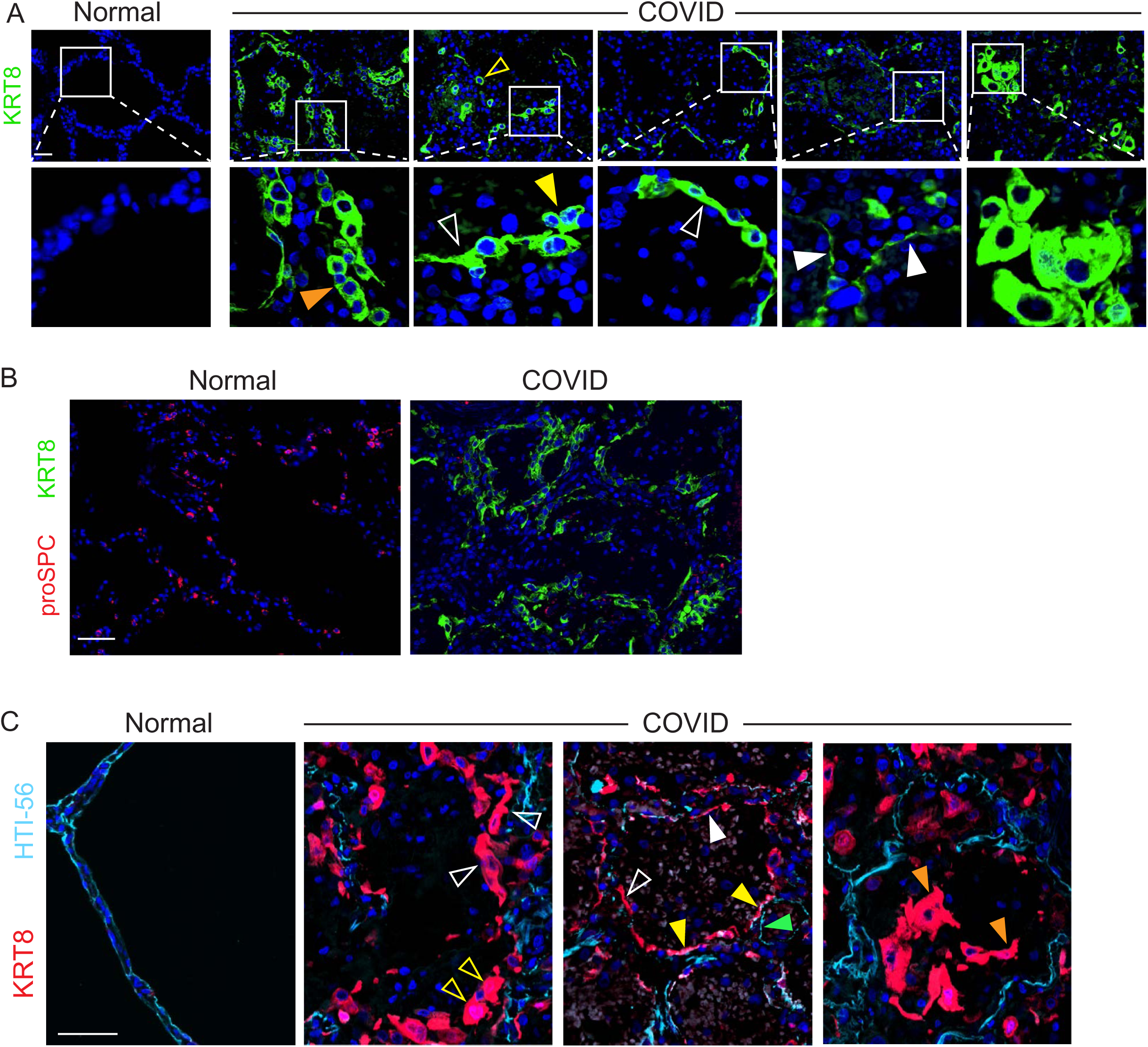
Epithelial Proliferation Yields Abundant Transitional Cells with Incomplete AEC1 Differentiation in COVID-19 ARDS. Immunostaining revealed A) KRT8^hi^ transitional cells that are sometimes cuboidal and isolated (open yellow arrowheads) or existing in pairs (closed yellow arrowheads) or hyperplastic (orange arrowhead), suggestive of recent cell division. Transitional cells were most often partially spread (open white arrowheads) and occasionally flat (closed white arrowheads). Scale bar = 50 µm. B) Transitional cells did not express proSPC and were abundant in areas devoid of SPC+ cells. Scale bar = 50 µm. C) Sections were stained for KRT8 and the AEC1 marker HTI-56. KRT8^hi^ transitional cells fill gaps denuded of AEC1s on alveolar septa. Some are cuboidal (open yellow arrowheads); most are partially spread (open white arrowheads). Occasional transitional cells approach a flat AEC1 morphology but do not express AEC1 markers (closed white arrowheads) with rare exception (closed yellow arrowheads). Flat cells that express AEC1 markers but not KRT8 (closed green arrowheads) were interpreted as native AEC1s that were not damaged during lung injury. Findings suggest ongoing organized, albeit incomplete, AEC1 differentiation. Rare cells display bizarre morphologies and/or had sloughed into the airspaces (orange arrowheads), consistent with haphazard regeneration. Scale bar = 50 µm. n=3.

### Incomplete Differentiation of Transitional Cells into Mature AEC1s in COVID-19 ARDS

Since progenitors successfully proliferated and assumed the transitional state, we hypothesized that incomplete differentiation of transitional cells into mature AEC1s may underlie the persistent barrier permeability, ongoing edema, and poor clinical outcomes in COVID-19. Supporting this hypothesis, although the transitional cells appeared to be in the process of differentiating into AEC1s, they were typically cuboidal or partially spread, only occasionally assuming the flat morphology of AEC1s (Figure 2A). To further elucidate the extent to which the transitional cells had differentiated into AEC1s, we costained lung sections for KRT8 and AEC1 markers (Figure 2C, Supplemental Figure 7). In most cases, transitional cells expanded along alveolar septa filling gaps denuded of AEC1s (Figure 2C). Again, some cells were cuboidal (open yellow arrowheads in Figure 2C), but most cells were partially spread (open white arrowheads in Figure 2C). However, some had a flat morphology (closed white arrowheads in Figure 2C) and occasional transitional cells expressed AEC1 markers (closed yellow arrowheads in Figure 2C). Flat cells expressing AEC1 markers but not KRT8 were interpreted as native AEC1s that withstood injury (closed green arrowheads in Figure 2C), although nascent AEC1s that downregulated KRT8 after differentiation cannot be excluded. Rarely, transitional cells assumed bizarre morphologies and/or had sloughed off the septa into the airspaces (closed orange arrowheads in Figure 2C). Although there was some variability between patients and across the tissue of each patient, the general histology, AEC1 damage, paucity of mature AEC2s, and abundance of transitional cells were present in all patients (Supplemental Figures 2, 6, 7, 8). The predominant organization of transitional cells in a monolayer on alveolar septa filling gaps denuded of AEC1s and displaying increasingly spread morphologies without AEC1 marker expression (Figure 2, Supplemental Figure 7) suggested that differentiation along a linear trajectory into mature AEC1s is ongoing but incomplete. Thus, the progressive edema, ongoing ventilator dependence and high mortality observed in COVID-19 ARDS may be due to incomplete differentiation of transitional cells into AEC1s.

### SARS-CoV-2 Was Not Detected in COVID-19 ARDS Autopsy Specimens

To assess for ongoing viral infection, which could contribute to the persistence of the transitional state, we examined the tissue for the presence of SARS-CoV-2. SARS-CoV-2 was not detected by immunostaining with an antibody against nucleocapsid protein (data not shown). Fluorescence *in situ* hybridization using a probe against SARS-CoV-2 failed to detect viral RNA, although staining with control probes suggested RNA degradation in the fixed tissue (data not shown). Viral inclusions were not observed, consistent with prior studies ^46^. Since SARS-CoV-2 infects AEC2s due to high expression of the viral receptor ACE2, and mature AEC2s were rare (Figure 1G, Supplemental Figure 2), we hypothesized that the apparent lack of ongoing viral infection may be due to a lack of ACE2 expression by transitional cells and AEC1s. Interrogation of publicly available scRNAseq datasets revealed that AEC2s downregulate ACE2 as they assume the transitional state and differentiate into AEC1s (Supplemental Figure 9). While it is difficult to make definitive conclusions based on negative data, the inability to detect SARS-CoV-2, absence of viral inclusions, and paucity of AECs that express the viral receptor argue against ongoing viral infection.

### Absence of Fibrosis in COVID-19 ARDS

In humans, persistence of the transitional state has previously been observed only in the setting of fibrosis, whether due to IPF or fibroproliferative ARDS, leading to speculation that this finding may be pathognomonic of fibrosis ^19, 20, 22, 24, 30, 31, 40, 41^. However, it is unknown whether transitional cells are inevitably associated with fibrosis in humans ^40, 41^. In mouse models of physiologic regeneration such as the LPS model and the pneumonectomy model, transitional cells appear transiently and then differentiate into AEC1s to restore normal alveolar architecture without fibrosis ^18, 21^. Therefore, we hypothesized that in early COVID-19 ARDS, transitional cells, which appear to be in the process of physiologic AEC1 differentiation (Figure 2C), exist without fibrosis. Indeed, we observed no excessive collagen deposition, myofibroblast accumulation, or architectural distortion in early COVID-19 ARDS, in contrast to IPF (Figure 3A,B). Moreover, whereas in IPF the transitional cells overlie fibroblastic foci (Figure 3C) and line honeycomb cysts ^22^, the transitional cells in COVID- 19 ARDS overlie structurally normal alveolar septa (Figure 3A Supplemental Figure 7). Finally, in early COVID-19 ARDS, nascent AEC1s appear to be emerging from transitional cells (Figure 2A,C, Supplemental Figure 7), whereas IPF is characterized by a paucity of AEC1s (Figure 3B).

**Figure 3.**
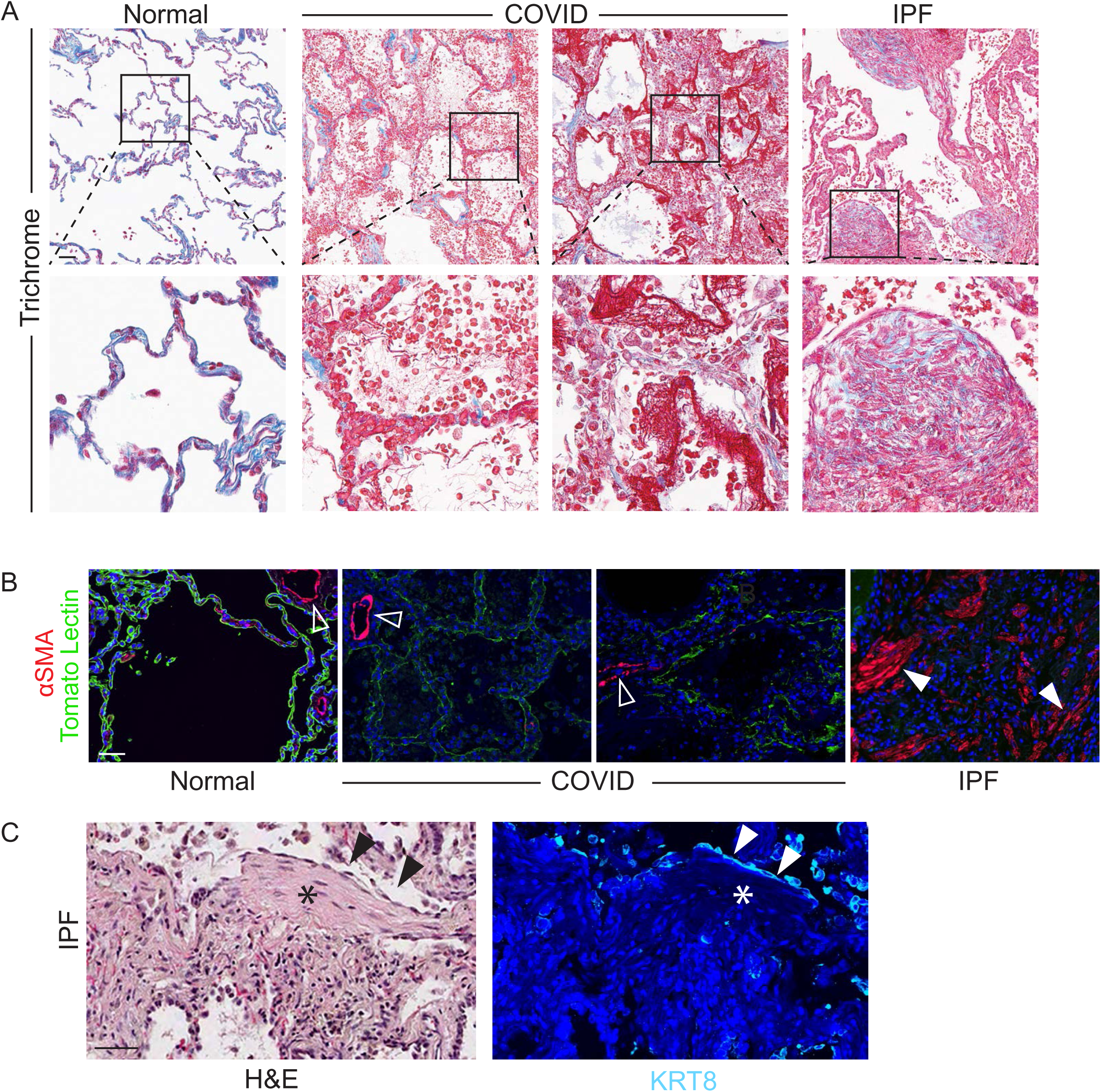
Absence of Fibrosis in Early COVID-19 ARDS. A) Trichrome highlighted (in blue) basement membranes and vascular adventitia in normal and COVID-19 lungs and collagen deposition with marked fibrosis in IPF. Scale bar = 50 µm. B) Immunostaining demonstrated abundant myofibroblasts in IPF but not COVID-19. Open arrowheads indicate smooth muscle cells; closed arrowheads indicate myofibroblasts. Scale bar = 50 µm. C) Serial sections were stained for H&E (left panel) or KRT8 (right panel). Transitional cells (black and white arrowheads) overlie fibroblastic focus (black and white asterisks). Scale bar = 50 µm. n=3.

### Non-COVID-19 ARDS Characterized by Accumulation of Transitional Cells Without Fibrosis

Although initial studies suggested high mortality rates for COVID-19 ARDS ^1^, recent evidence indicates that clinical outcomes of COVID-19 ARDS may be similar to those of ARDS from other etiologies ^47^. Since the histology of COVID-19 ARDS is identical to that of ARDS of other etiologies [classic DAD (Figures 1A-D, Supplemental Figure 8) ^26–28, 48^], we hypothesized that the state of regeneration observed here is common to fatal early ARDS regardless of etiology. To address this hypothesis, we immunostained post-mortem lung tissue from patients who died within 14 days of ARDS of etiologies other than COVID-19. These patients had acute DAD (Supplemental Figure 10) with AEC1 damage (Supplemental Figure 11A), AEC2 proliferation (Supplemental Figure 11B,12), abundant transitional cells (Supplemental Figure 11C), and incomplete AEC1 differentiation (Supplemental Figure 11D) in the lungs of patients who died of ARDS of other etiologies. However, in contrast to COVID-19 ARDS lungs, which were largely devoid of mature AEC2s, mature AEC2s were typically present throughout the non-COVID-19 ARDS lungs (Supplemental Figures 11B, 12). In addition, occasional cells were proSPC+ KRT8^hi^ (Supplemental Figure 12). The non-COVID-19 ARDS lungs were without fibrosis (Supplemental Figure 13)

### Transitional Cells are Senescent in IPF but not in ARDS

It is unknown why in mouse models of physiologic alveolar regeneration and early human ARDS, transitional cells appear capable of differentiating into AEC1s with restoration normal alveolar architecture, whereas in IPF, transitional cells persist with a paucity of AEC1s and fibrosis ensues. To confirm that the transcriptome of transitional cells is highly conserved across diverse mouse models of physiologic alveolar regeneration, human ARDS, and human IPF, we assessed expression of transitional state markers by immunostaining and interrogation of existing scRNAseq datasets. Transitional cells in the LPS and PNX mouse models of physiologic alveolar regeneration and human IPF shared the unique gene expression signature of the transitional state (Figure 4A). That the transcriptome of the transitional state was highly conserved across two mouse models of physiologic alveolar regeneration and IPF underscores the enigma of the vastly divergent pathologic outcomes. During physiologic regeneration and likely in early human ARDS (Figure 2C, Supplemental Figure 7), transitional cells emerge as proliferating AEC2s exit the cell cycle and express markers of cell cycle arrest, but ultimately differentiate into AEC1s ^18, 21^, whereas AECs in IPF have previously been identified as senescent ^30–35, 40^. Therefore, we speculated that the critical difference underlying the divergent outcomes between physiologic regeneration, early human ARDS, and fibrosis may be that while in physiologic regeneration and early human ARDS, transitional cells assume a transient state of cell cycle arrest as proliferating AEC2s exit the cell cycle in anticipation of terminal differentiation, over time in the context of ongoing or repetitive injury, they evolve into a permanent state of cell cycle arrest, or senescence, and fibrosis ensues. To confirm that transitional cells in fibrosis but not mouse models or human ARDS are senescent, we assessed expression of general markers of cell cycle arrest, p21 (CDKN1A), p15 (CDKN2B), p53 (TP53), and cyclin D1 (CCND1) and a more specific marker of senescence, p16 (CDKN2A) ^49^, in mouse models, early human ARDS, and human IPF. We found that general markers of cell cycle arrest are expressed in all three conditions (Figure 4B,C), but p16 is expressed only in IPF (Figure 4D,E, Supplemental Figure 14). These findings were conserved across the patients studied. Notably, both general markers of cell cycle arrest and p16 are expressed in transitional cells in the fibrosis associated with fibroproliferative ARDS ^40^. Taken together, these data suggest that in physiologic regeneration in mice and early human ARDS, proliferating AEC2s undergo cell cycle exit and adopt the transitional state *transiently* before differentiating into AEC1s to restore normal alveolar architecture, whereas, in fibrosis due to IPF or fibroproliferative ARDS, transitional AECs eventually become *permanently* arrested, or senescent, losing capacity for an AEC1 fate and begetting fibrosis (Supplemental Figure 15).

**Figure 4.**
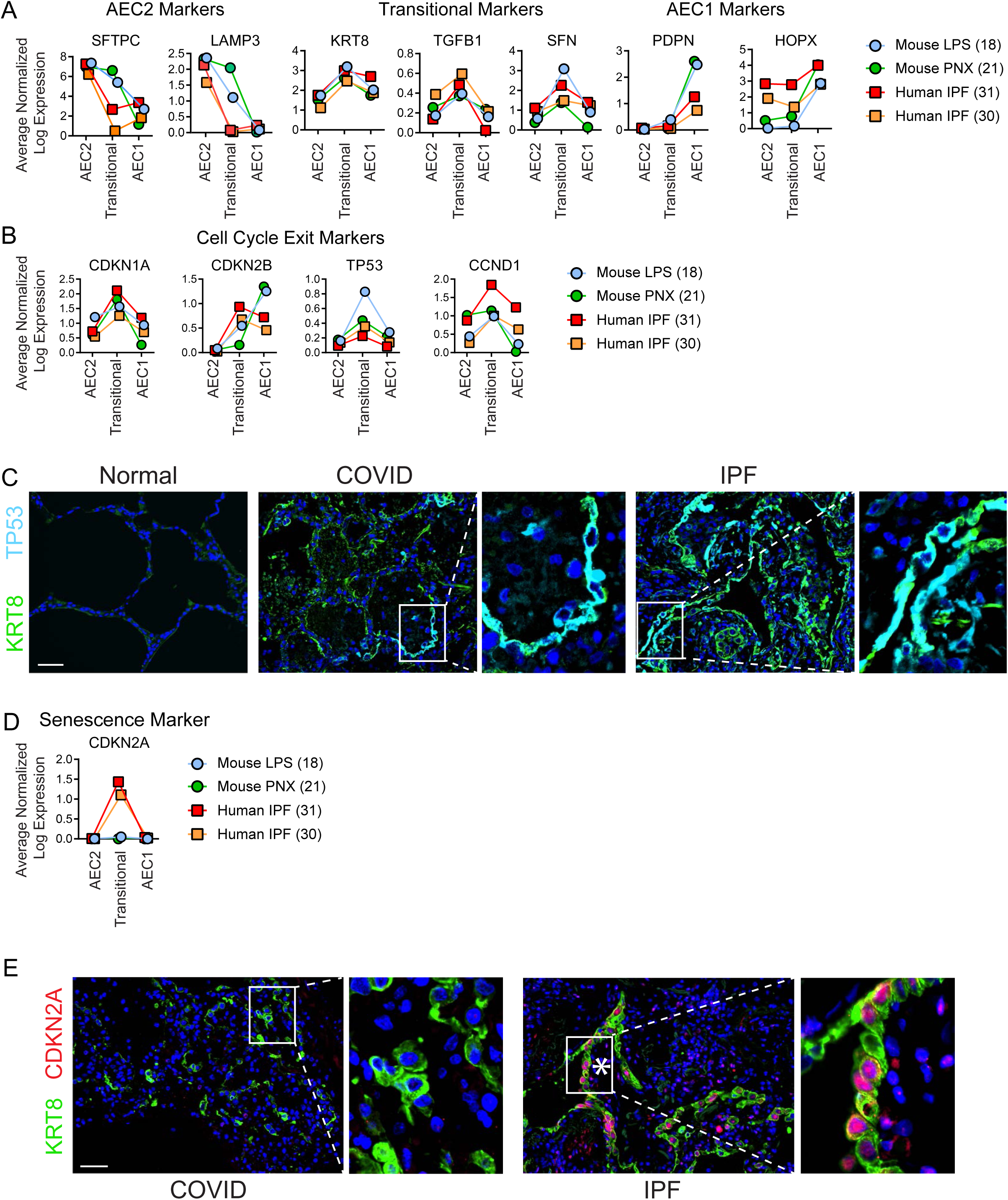
Transitional Cells in Mouse Models of Physiologic Regeneration, COVID-19 ARDS, and IPF Exist in a State of Cell Cycle Exit but only in IPF are Senescent. A,B,D) Single cell RNA sequencing datasets from two mouse model of physiologic regeneration, LPS ^18^ and pneumonectomy (PNX) ^21^ and human IPF ^30, 31^ were interrogated. C,E) Lung sections were immunostained. A) As AEC2s assume the transitional state, they downregulate AEC2 markers and upregulate transitional markers which are conserved in mouse models of physiologic regeneration and human IPF. AEC1s express low levels of transitional markers and high levels of AEC1 markers. In mouse models of physiologic regeneration, COVID-19 ARDS, and IPF, transitional cells express general markers of cell cycle exit (B,C), but only in IPF do they express the highly specific marker of senescence CDKN2A/p16 (D,E). C,E) Asterisks indicate fibroblastic foci. Scale bars = 50 µm. n=3.

## Discussion

Herein, we performed a comprehensive examination of epithelial injury and regeneration in early, fatal ARDS due to COVID-19 and other etiologies, with comparison to IPF. The early ARDS lungs were characterized by extensive epithelial damage and a regenerative response in which AEC2s (or other progenitors) proliferated and assumed the transitional state. Residual mature AEC2s were rare in the COVID-19 but not in the non-COVID-19 lungs. Some transitional cells were hyperplastic and cuboidal. However, most displayed a partially spread morphology, filling gaps on alveolar septa denuded of AEC1s, suggesting they were in the process of AEC1 differentiation. The transitional cells occasionally assumed a flat AEC1 morphology and rarely expressed AEC1 markers, suggesting that AEC1 differentiation was ongoing but incomplete. Rare transitional cells appeared to have sloughed off into the airspaces. Similar to mouse models of physiologic regeneration and in contrast to IPF and fibroproliferative ARDS, transitional cells existed on structurally normal alveolar septa without fibrosis and expressed markers of cell cycle exit but not senescence. Taken together, these data suggest that transitional cells arise in early human ARDS without fibrosis and that AEC1 differentiation from the transitional state with restoration of normal alveolar architecture and barrier integrity is ongoing but incomplete in early fatal ARDS, whereas impaired AEC1 differentiation and ensuing fibrosis are irreversible in fibroproliferative ARDS and IPF.

In ARDS, injury to the alveolar epithelial barrier results in noncardiogenic pulmonary edema, which causes severe hypoxemia frequently necessitating mechanical ventilation and leading to death ^6^. Conversely, epithelial repair is associated with clinical recovery and survival ^10, 11^. However, whether poor clinical outcomes in ARDS may be linked to ineffectual epithelial regeneration is unknown. Our study revealed that progenitors proliferate and assume the transitional state but that incomplete AEC1 differentiation from the transitional state likely explains the persistent barrier permeability, ongoing pulmonary edema, ventilator dependence, and death in these patients.

Future research should explore whether specific factors actively delay AEC1 differentiation from the transitional state. An intriguing possibility would be that infection of transitional cells by SARS-CoV-2 maintains cells in the transitional state. We did not detect viral nucleocapsid protein or viral inclusions. Although negative results are difficult to interpret, the absence of active viral infection would be consistent with prior work demonstrating that COVID-19 patients rarely have live virus after Day 8 of infection despite amplification of viral RNA by PCR ^29, 40, 50, 51^. Moreover, it is unlikely that transitional cells become infected by SARS-CoV-2 since they do not express its receptor, ACE2 (Supplemental Figure 9) ^20, 22^. In fact, given the dramatic paucity of mature AEC2s in COVID-19 ARDS (Figure 1G, Supplemental Figure. 2), lack of persistent SARS-CoV-2 infection may be due to depletion of the AEC2 reservoir. Persistence of the transitional state, also observed in non-COVID-19 ARDS, is likely due to signals from within the transitional cells or from the injured lung milieu. Our previous work suggested that TGFβ maintains the transitional state ^18^; other transcriptional pathways differentially activated in the transitional and AEC1 states such as Sox4 ^18–21^ merit further investigation. Additionally, unchecked inflammation, which is known to maintain the transitional state ^24^ and contribute to poor clinical outcomes in ARDS ^52^ may delay AEC1 differentiation. The nongradual nature of differentiation suggests that activation of some process is necessary to induce transitional cells to differentiate into AEC1s. Elucidation of the cellular and molecular mechanisms regulating AEC1 differentiation may ultimately lead to novel therapies to accelerate AEC1 differentiation, barrier restitution, liberation from the ventilator, and survival in ARDS.

Transitional cells had previously been identified in humans only in the setting of fibrosis, whether in the context of IPF or fibroproliferative ARDS ^19, 20, 22, 24, 31, 40, 41^. In fact, it has been proposed that persistence of the transitional state with impaired AEC1 differentiation is the specific regenerative defect driving the pathogenesis of fibrosis and that the presence of transitional cells on lung biopsy may serve as a biomarker of fibrosis in prolonged ARDS, indicating to the clinician that lung transplant may be the only viable therapeutic option ^19–22, 40^. However, it was unknown whether transitional cells are inevitably associated with fibrosis in humans. During physiologic regeneration in mice, proliferating AEC2s exit the cell cycle and transiently assume the transitional state before differentiating into AEC1s to restore normal alveolar architecture and function without fibrosis^18, 21^. Here, we demonstrate for the first time that transitional cells arise in early human ARDS *without fibrosis*. Although AEC1 differentiation from the transitional state was incomplete in early ARDS, the appearance of cuboidal, partially spread, and flat transitional cells organized in a monolayer on structurally normal alveolar septa without fibrosis suggests that these cells may retain the capacity for physiologic AEC1 differentiation with restoration of normal alveolar architecture and barrier integrity, as occurs in mouse models of physiologic regeneration ^18, 21^. This provides pathophysiologic rationale that clinical recovery may be possible and justifies ongoing aggressive supportive care, including extracorporeal membrane oxygenation (ECMO), to allow time for complete regeneration. Of course, had the patients studied here survived and suffered additional insults to the epithelium from prolonged mechanical ventilation and hospital acquired pneumonias, the transitional cells may have eventually lost the capacity for AEC1 differentiation and become senescent with ensuing fibrosis. However, the novel finding of the present study is that transitional cells arise early during regeneration in ARDS in the absence of fibrosis, at which time they appear to retain capacity for AEC1 differentiation and restoration of normal lung structure and function.

By comparing and contrasting the transitional state in mouse models of physiologic regeneration, early human ARDS, and human IPF, our study provides insight into the fundamental question of why injury sometimes resolves with physiologic regeneration and other times leads to fibrosis. We confirmed that the transcriptomes of the transitional state in two mouse models of physiologic regeneration and human IPF are highly conserved. Taken together, these findings raise the pivotal question of why transitional cells may maintain the capacity to differentiate into AEC1s and restore normal alveolar structure in mouse models of physiologic regeneration and early human ARDS but persist in the transitional state, leading to fibrosis, in IPF and fibroproliferative ARDS. We discovered that while transitional cells in physiologic regeneration in mice, early human ARDS, and IPF all express markers of cell cycle exit, only in IPF do they express a specific marker of senescence. Therefore, we propose the novel paradigm that in mouse models of physiologic regeneration and early human ARDS, proliferating AEC2s exit the cell cycle and *transiently* adopt the transitional state but retain the capacity to differentiate into AEC1s, restoring normal alveolar architecture without fibrosis, whereas in IPF and fibroproliferative ARDS, transitional AECs evolve into a *permanent* state of cell cycle arrest, or senescence, losing capacity for an AEC1 fate and promoting fibrosis (Supplemental Figure 15). Additional investigation will be necessary to confirm this hypothesis and to determine whether senescent transitional cells actually cause fibrosis and the underlying mechanisms. Since transitional cells lie in close spatial proximity to fibroblasts in the fibroblastic foci (Figure 4C) ^22, 30^ and express profibrotic genes including TGFβ (Figure 4A) ^18, 30, 31^, we further speculate that transitional cells may promote fibrosis by directly stimulating fibroblasts to become myofibroblasts and deposit matrix. If evolution from a transient state of cell cycle exit into a permanent state of senescence proves to be the critical switch that irreversibly diverts physiologic regeneration towards fibrosis, elucidation of the mechanisms by which transitional cells become senescent and by which senescent transitional cells activate fibroblasts may lead to the development of novel therapies to promote physiologic regeneration and prevent fibrosis in IPF and fibroproliferative ARDS.

This comprehensive immunostaining of DAD lungs for AEC2, transitional cell, and AEC1 markers yielded some unanticipated discoveries that are clinically relevant and may shift paradigms in our basic understanding of the pathology of DAD. We found that both sloughed epithelial cells long believed to be AEC1s and some hyperplastic epithelial cells long believed to be AEC2s are actually transitional cells. These discoveries not only suggest a longstanding misidentification of epithelial cell state in DAD but provide additional insight into clinical outcomes. The dramatic paucity of mature, surfactant-producing AEC2s in COVID-19, whether due to cytopathic viral infection or acquisition of the SPC- negative transitional state, likely contributes to surfactant deficiency. Surfactant deficiency causes atelectasis, which exacerbates hypoxemia and poor lung compliance and leads to compensatory alveolar overdistension and ventilator- induced lung injury (VILI) ^6^. The dramatic downregulation of surfactant demonstrated here may be one reason why corticosteroids, which stimulate surfactant production ^53^, have clinical efficacy in COVID-19 ^54^. That many desquamated and hyperplastic epithelial cells long believed to be AEC1s and AEC2s, respectively, are actually transitional cells suggests that the histologic entity known as “acute DAD” represents a significantly more advanced state of regeneration than previously recognized. The acute injury phase of DAD probably occurs within the first few hours to days of clinically manifest ARDS and is unlikely to be captured on autopsy specimens.

Our study has several limitations. First, only 6 ARDS patients were studied. However, the findings were highly uniform across patients with ARDS of various etiologies, suggesting generalizability. Since ARDS will remain a prevalent and devastating cause of morbidity and mortality long beyond the COVID-19 pandemic ^6^, the findings presented here will have enduring impact. Findings were also highly conserved across the IPF patients studied. Second, all COVID-19 ARDS patients died after mechanical ventilation; however, the histology of the non-COVID-19 ARDS patients that were not ventilated was identical. Third, although a failure of barrier restitution due to incomplete AEC1 differentiation likely contributes to poor clinical outcomes, causality has not been established. The patients studied here died with refractory acute respiratory failure due to noncardiogenic pulmonary edema, and the histologic appearance of transitional cells (Figure 2, Supplemental Figures 4,7) certainly does not seem compatible with barrier integrity. Fourth, as mentioned, we have not excluded the possibility that some transitional cells arise from progenitors other than AEC2s ^15–17^, particularly in COVID-19 ARDS given the paucity of AEC2s. Fifth, although we speculate that transitional cells retain the ability to differentiate into AEC1s in early human ARDS and lose the capacity for AEC1 differentiation in fibroproliferative ARDS and IPF, the potential fates of these cells are unknown. Without the ability to acquire serial tissue samples, intervene, and perform lineage tracing, it is difficult to definitively establish causality and cell fate in observational human studies. Interpretation of cell fate dynamics remains speculative. Moreover, while p16 is considered specific for senescence, distinguishing senescence from transient cell cycle arrest is not straightforward ^49^. Finally, the notion that evolution from a transient state of cell cycle arrest to a truly senescent state is the critical defect underlying the irreversible switch from physiologic regeneration towards fibrosis has not been proven.

In conclusion, the findings presented here suggest that in early fatal human ARDS, physiologic AEC1 differentiation from transitional cells is incomplete, thus underlying prolonged barrier permeability and respiratory failure, but as in physiologic regeneration in mice, is ongoing without fibrosis. These findings establish a foundation for future mechanistic studies to dissect the molecular mechanisms by which transitional cells can differentiate into AEC1s during physiologic regeneration and early ARDS and may lose capacity for an AEC1 fate and/or promote fibrogenesis in fibroproliferative ARDS and fibrosis. Such mechanistic studies may ultimately lead to novel therapies to promote AEC1 differentiation, thus accelerating restoration of barrier integrity, clearance of edema fluid, liberation from the ventilator, and survival in severe ARDS and prevent fibrosis in fibroproliferative ARDS and IPF.

## Acknowledgments

Acquisition of tissue: Rachel Dyal. Thoughtful discussions: Bethany Moore, Marc Peters-Golden, Elizabeth Redente, Michael Matthay, Michael Beers, and Guy Zimmerman.

## Author Contributions

Conception and design: RLZ. Data acquisition and analysis: CT, MA, NV, SH, KAR, FW, CF. Interpretation of Data: CT, CF, RLZ. Drafting or revising the manuscript: CT, RLZ. Final approval: All authors. RLZ is the guarantor of this work and, as such, had full access to all of the data in the study and takes responsibility for the integrity of the data and the accuracy of the data analysis.

---

**Supplemental Figure 1.**
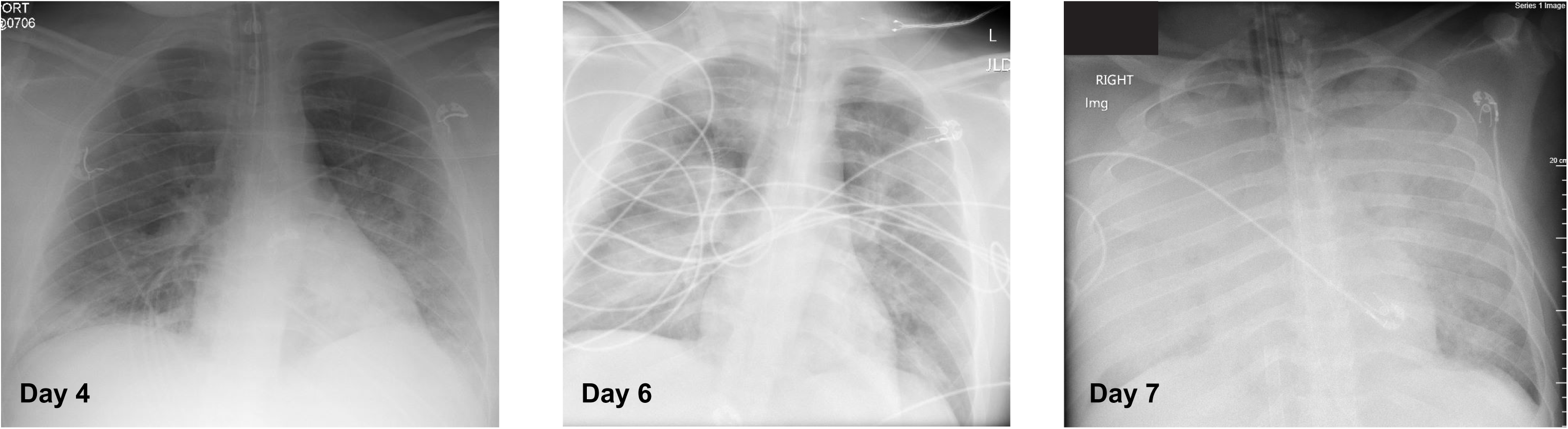

**Supplemental Figure 2.**
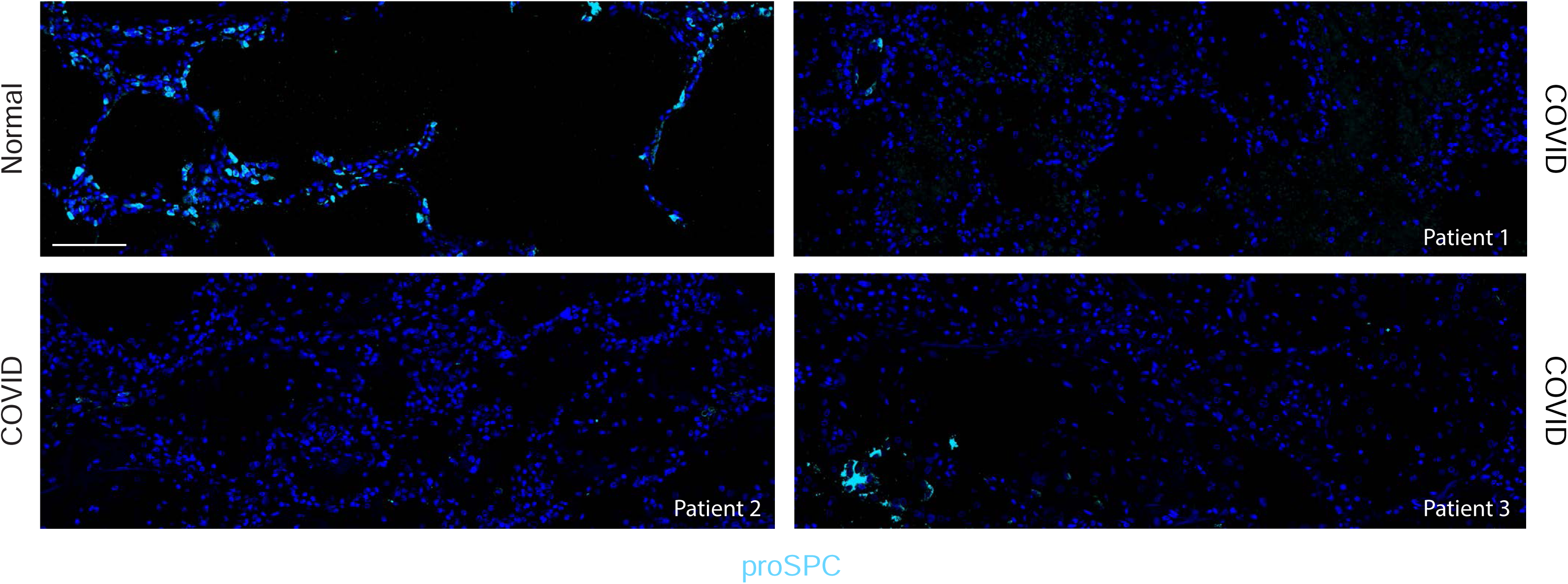

**Supplemental Figure 3.**
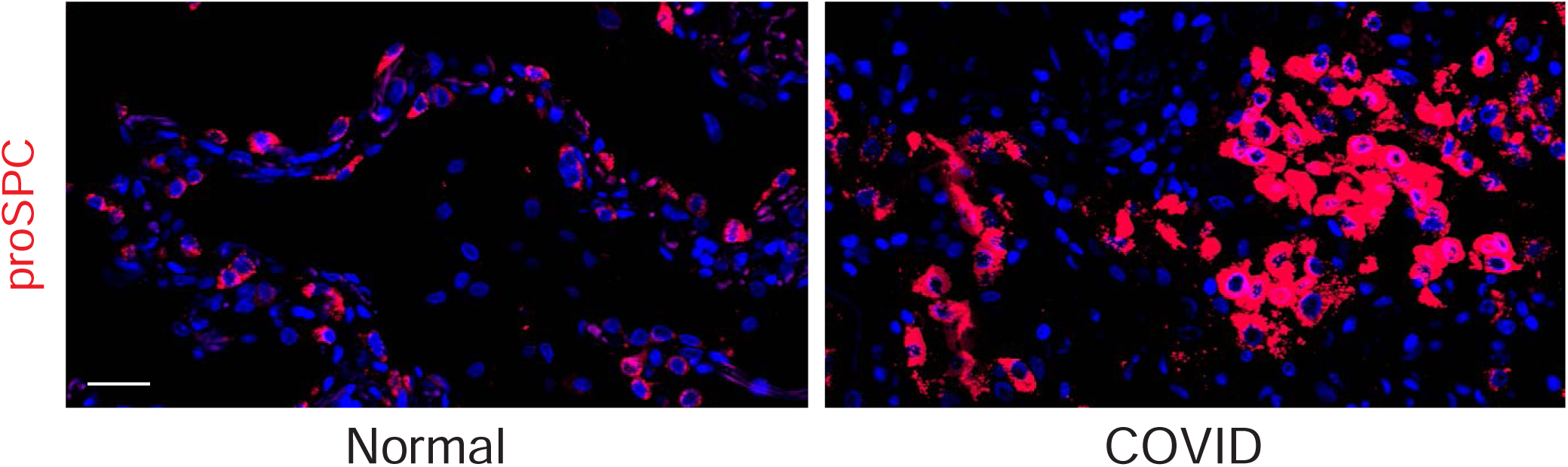

**Supplemental Figure 4.**
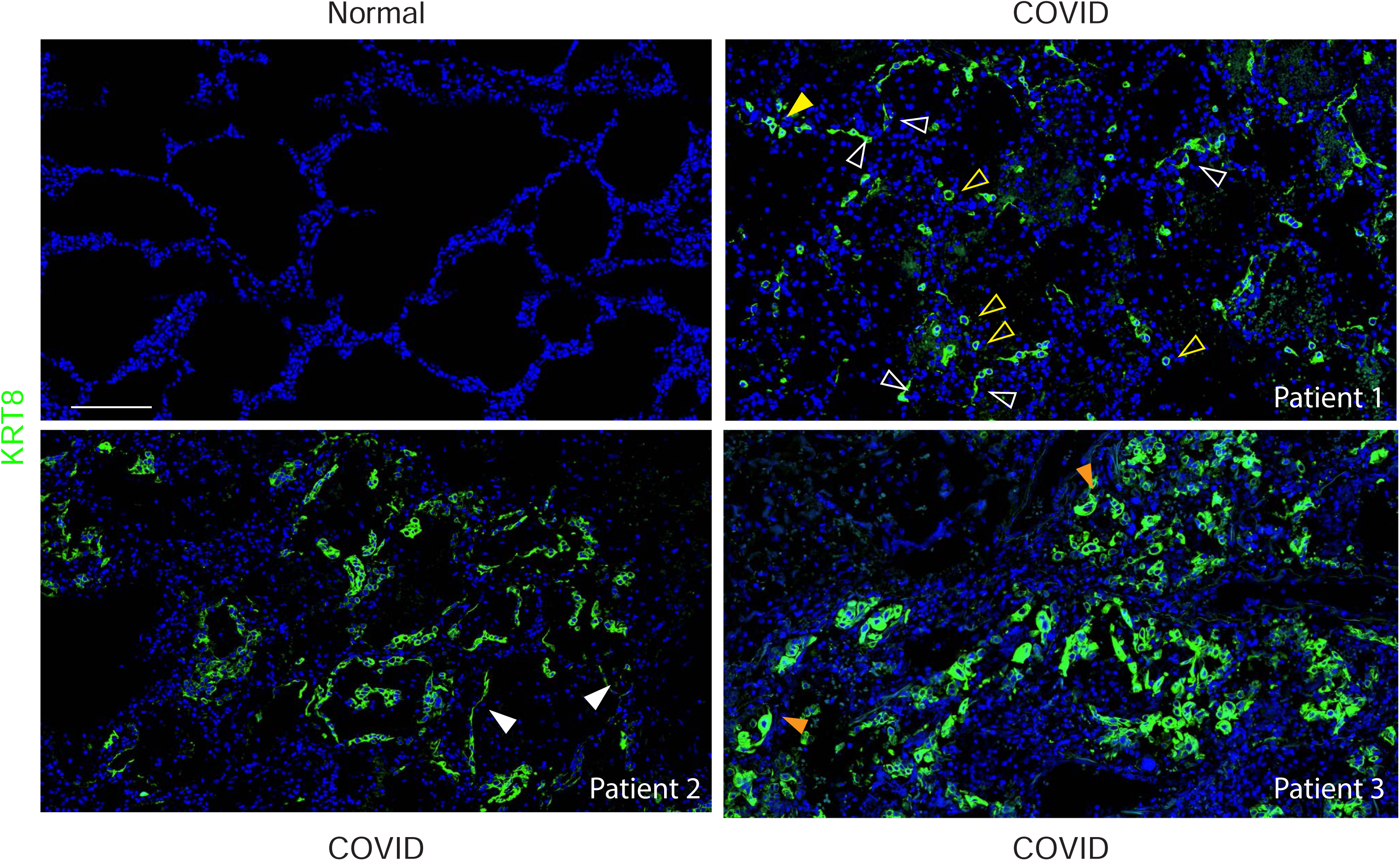

**Supplemental Figure 5.**
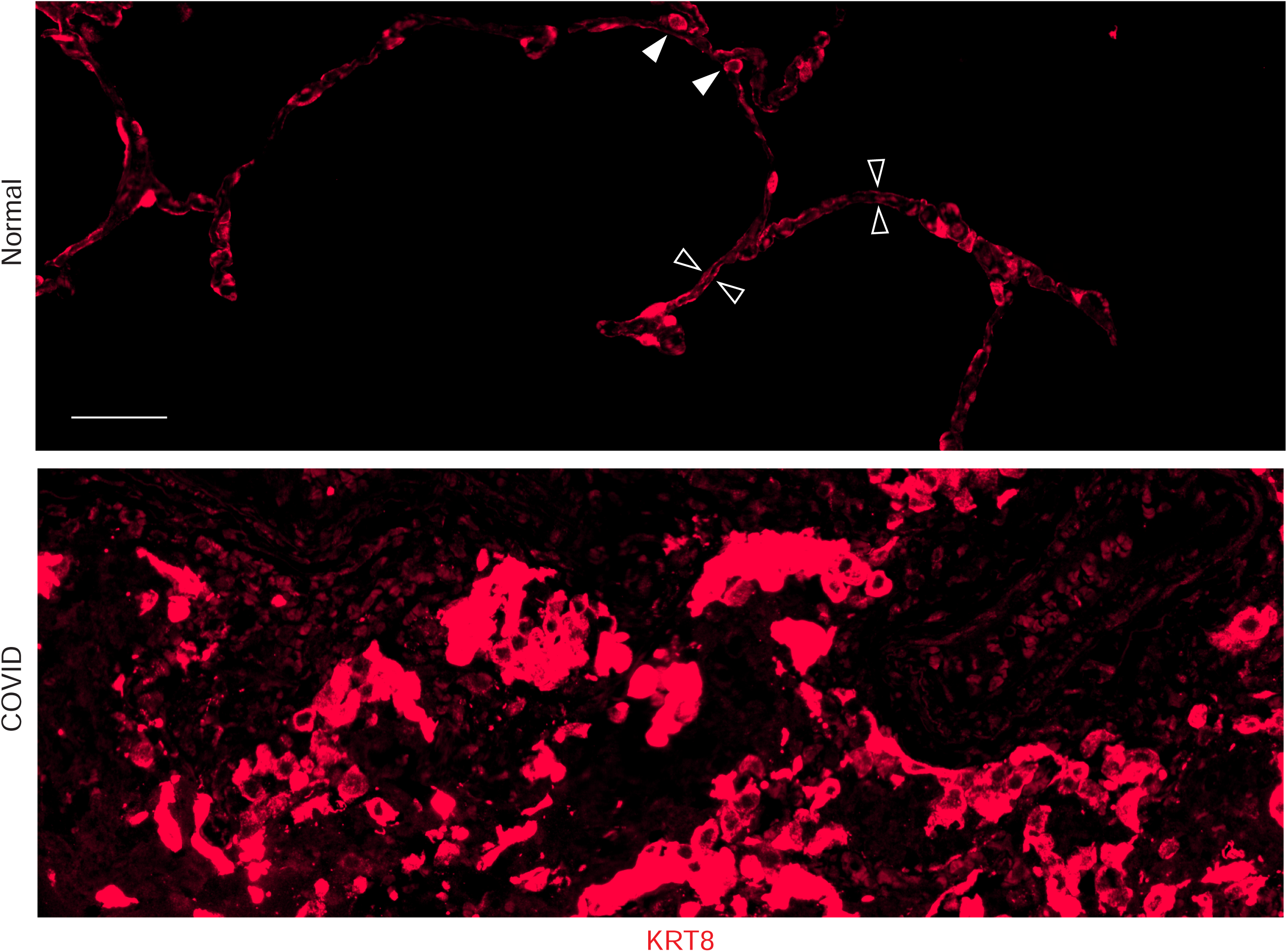

**Supplemental Figure 6.**
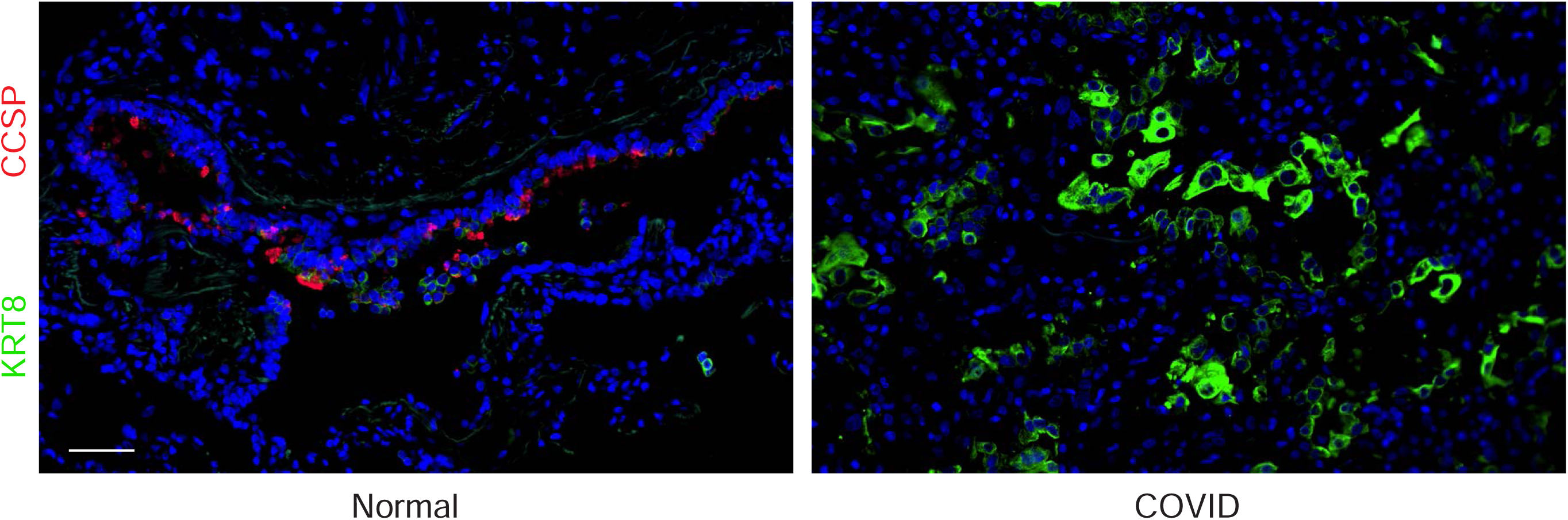

**Supplemental Figure 7.**
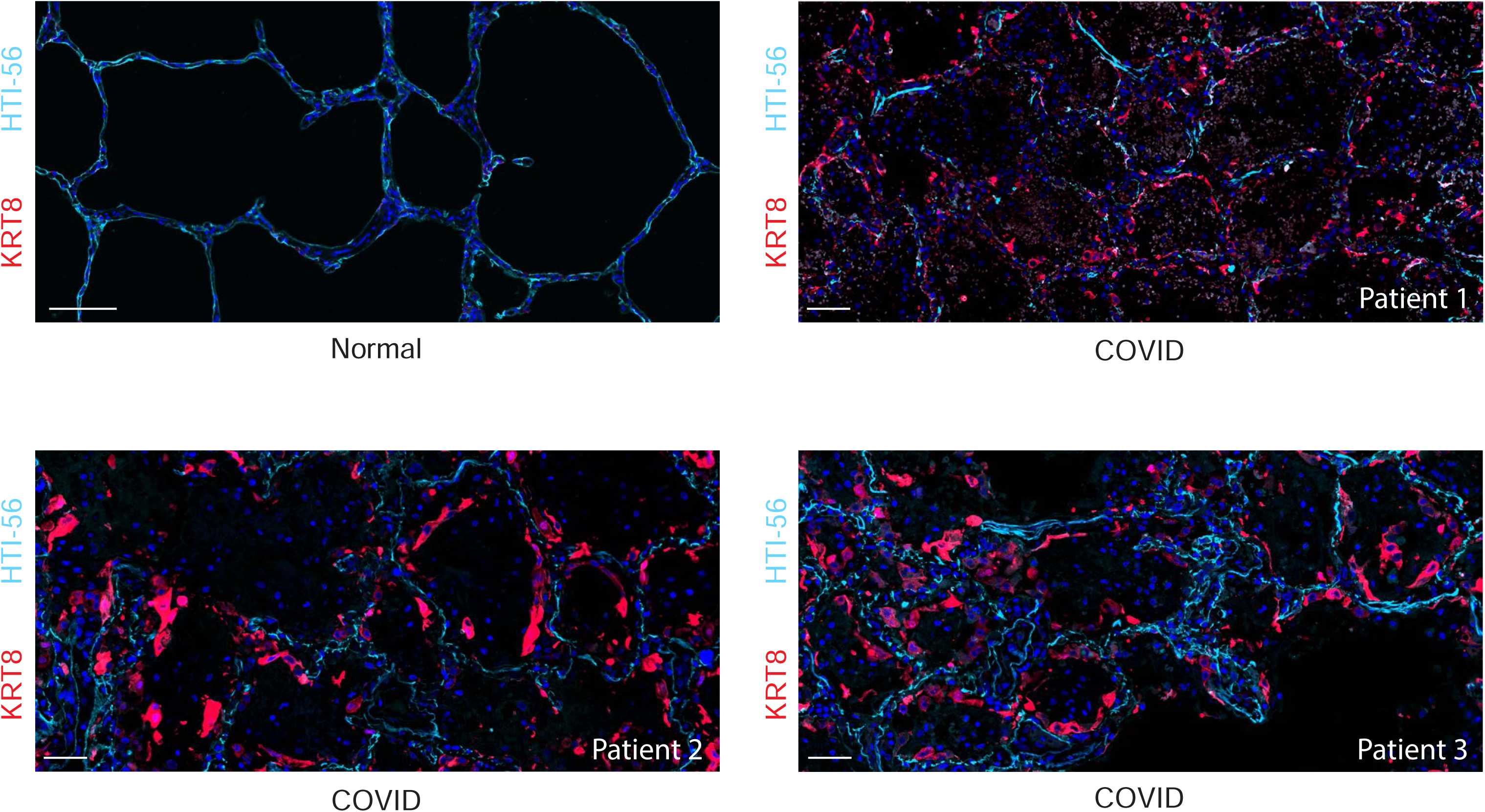

**Supplemental Figure 8.**
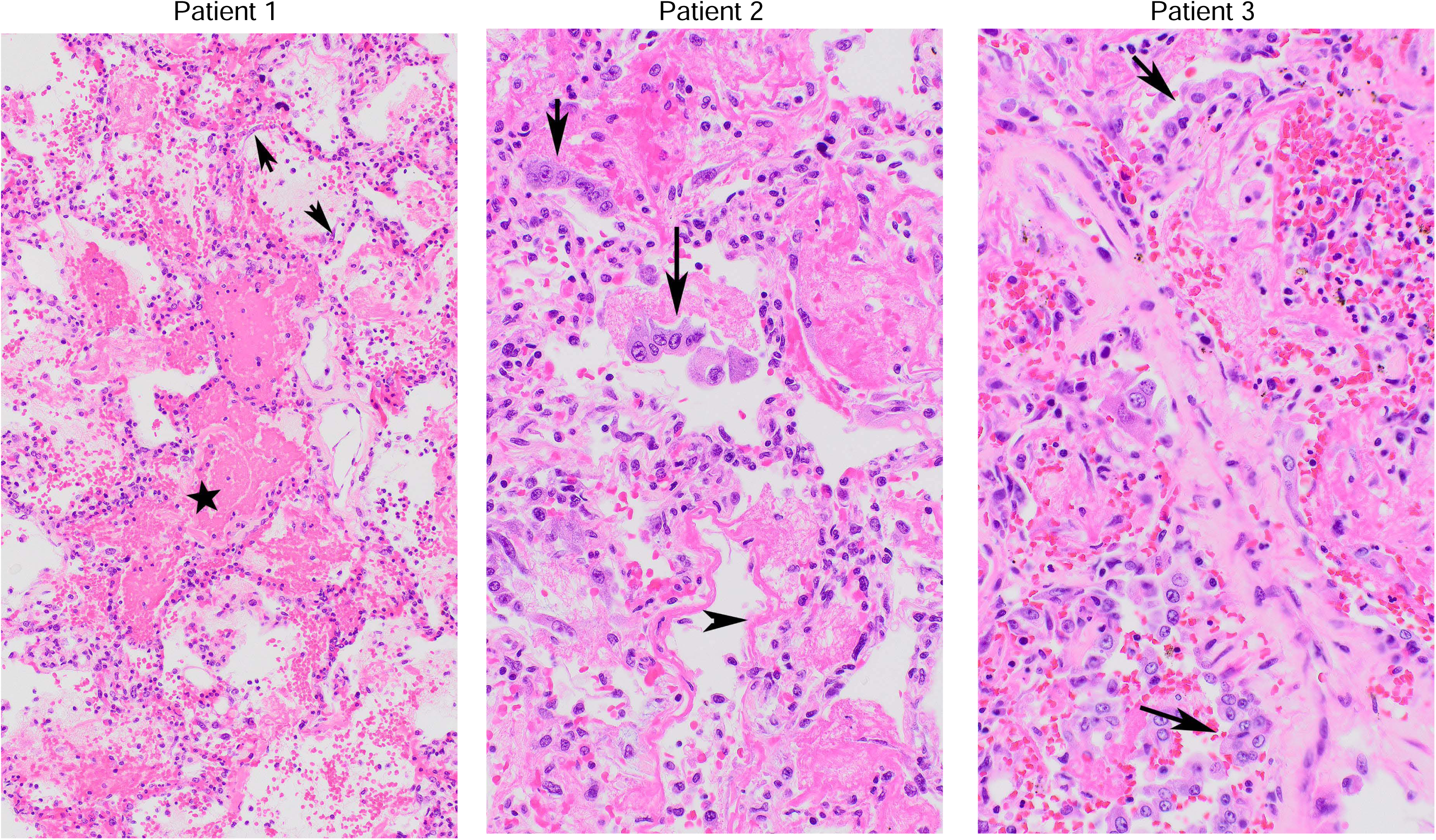

**Supplemental Figure 9.**
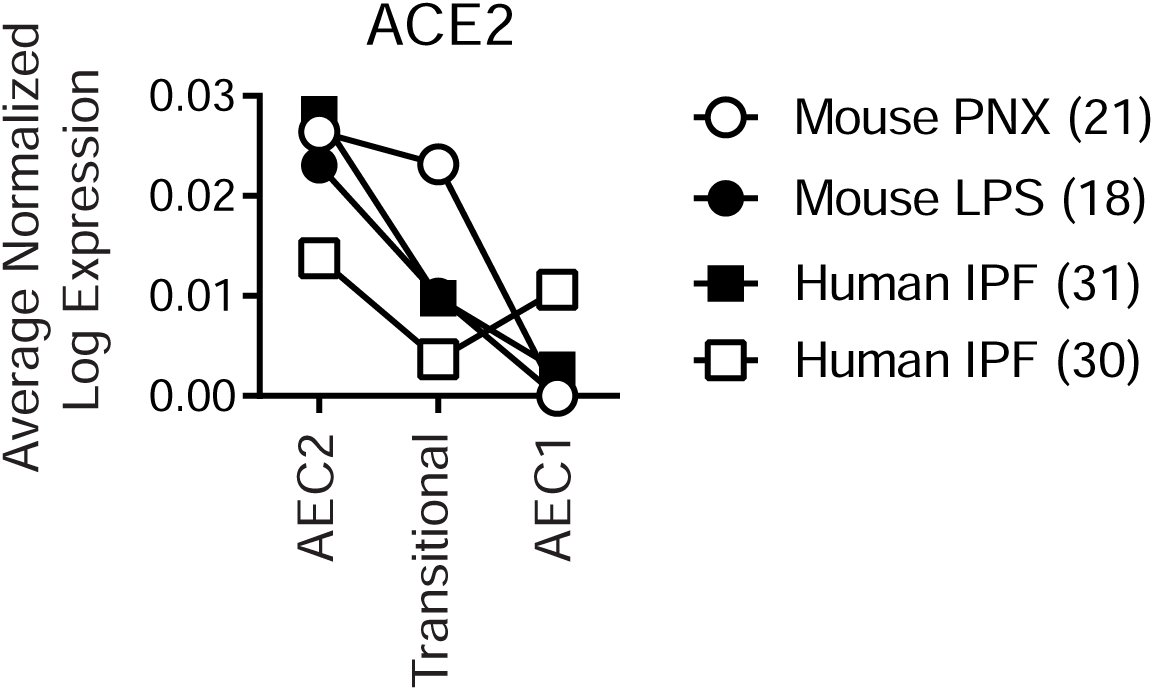

**Supplemental Figure 10.**
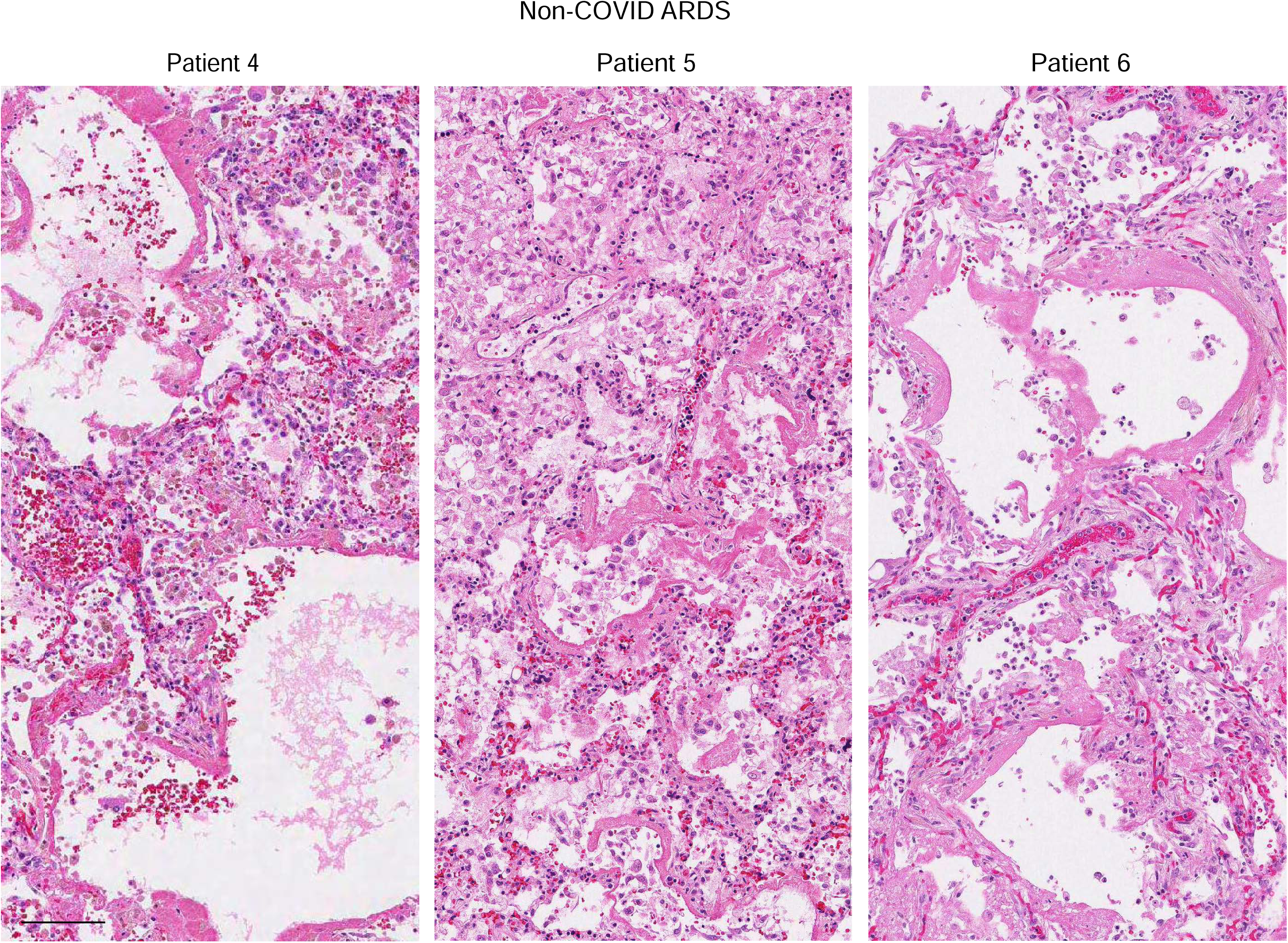

**Supplemental Figure 11.**
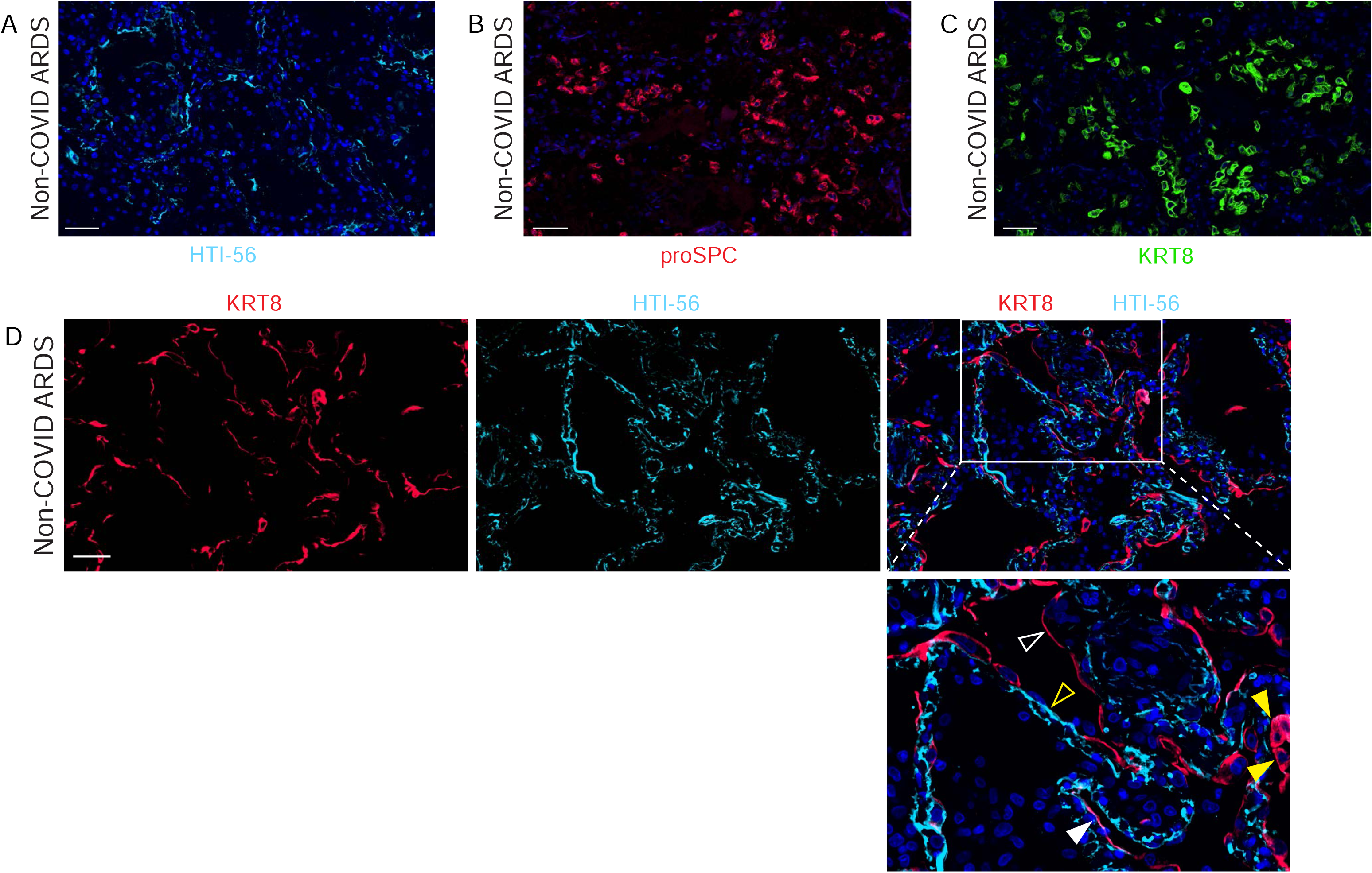

**Supplemental Figure 12.**
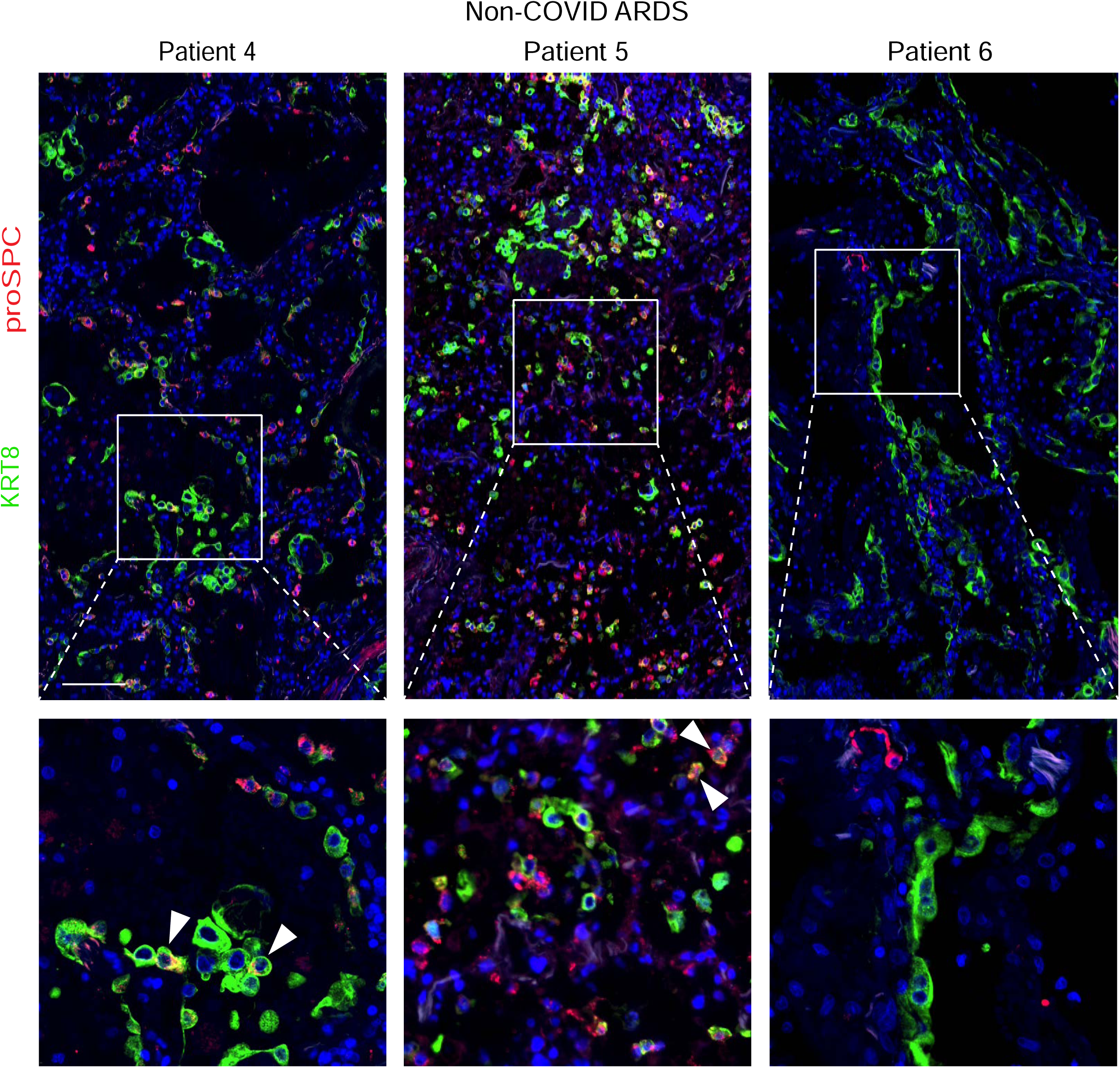

**Supplemental Figure 13.**
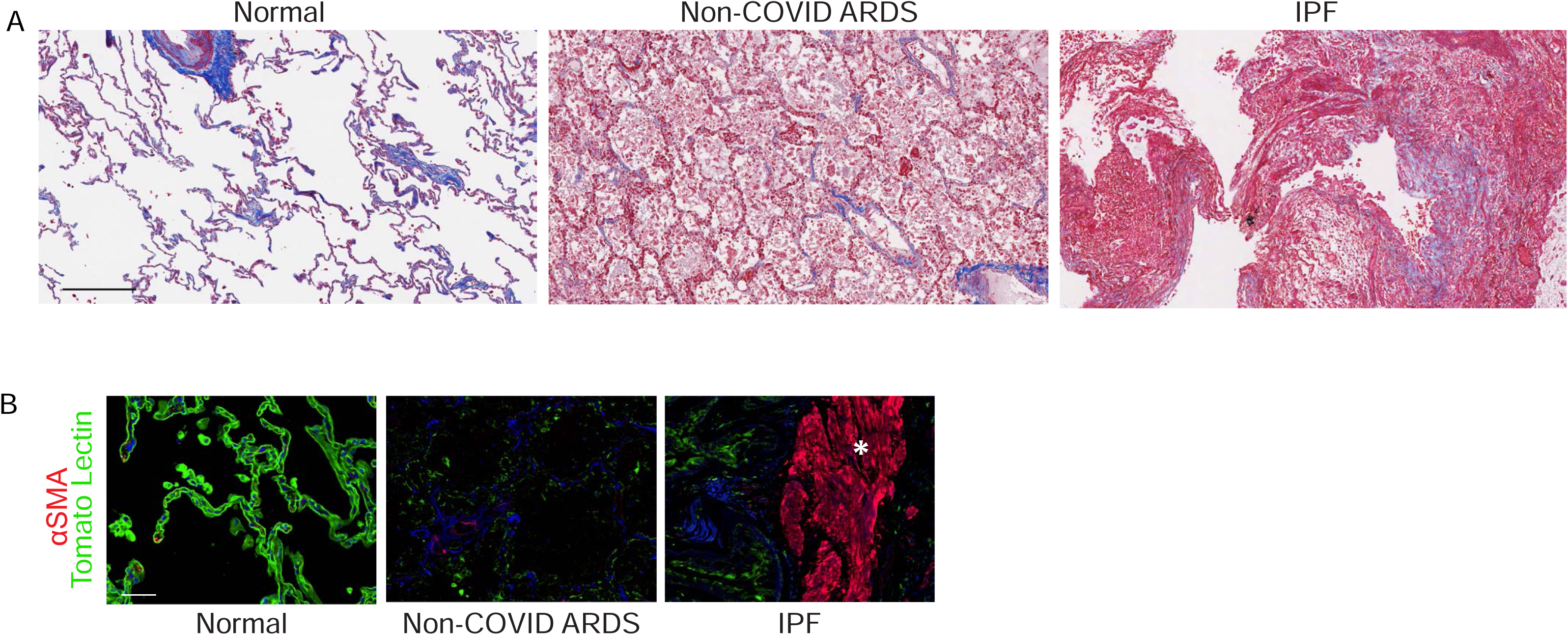

**Supplemental Figure 14.**
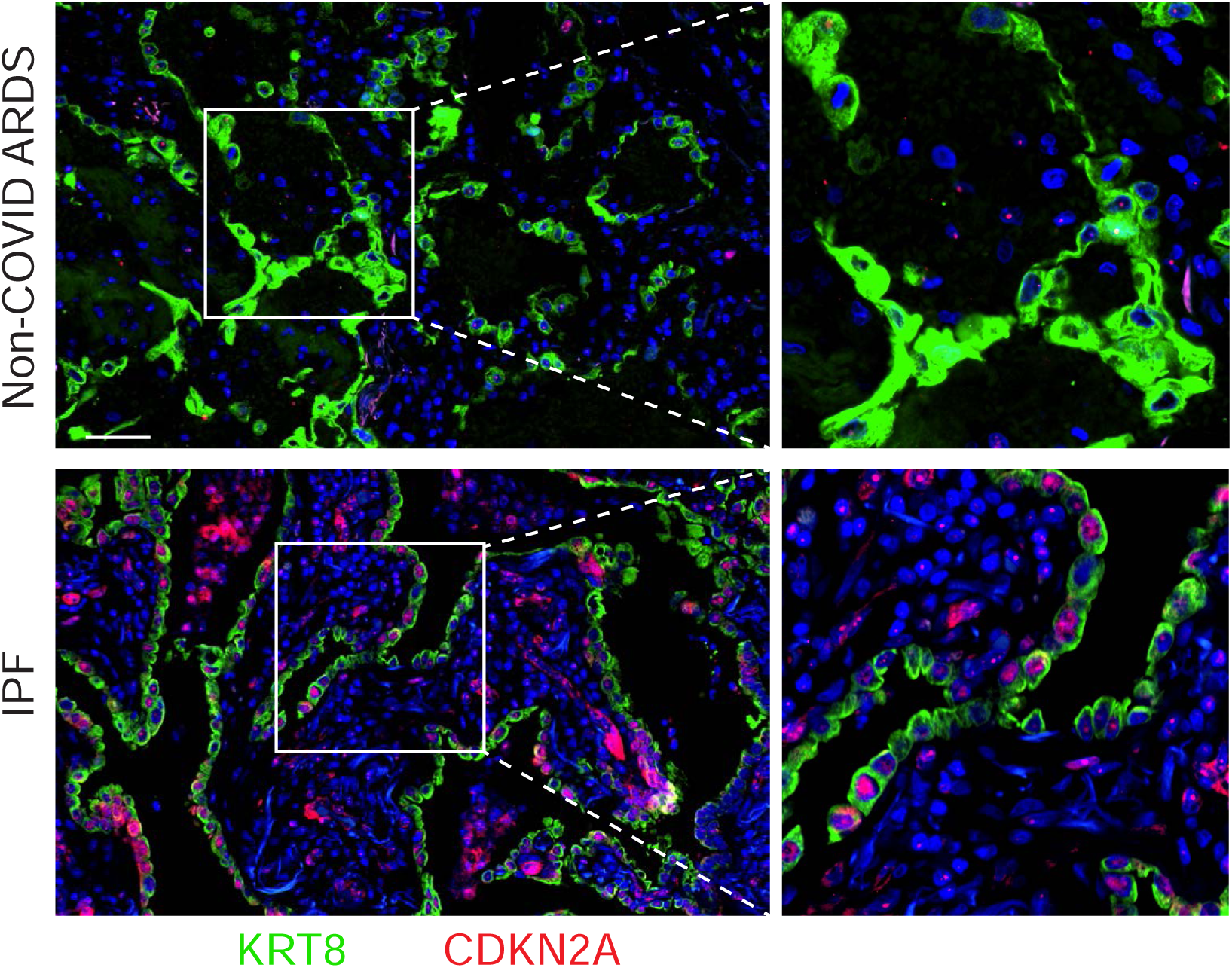

**Supplemental Figure 15.**
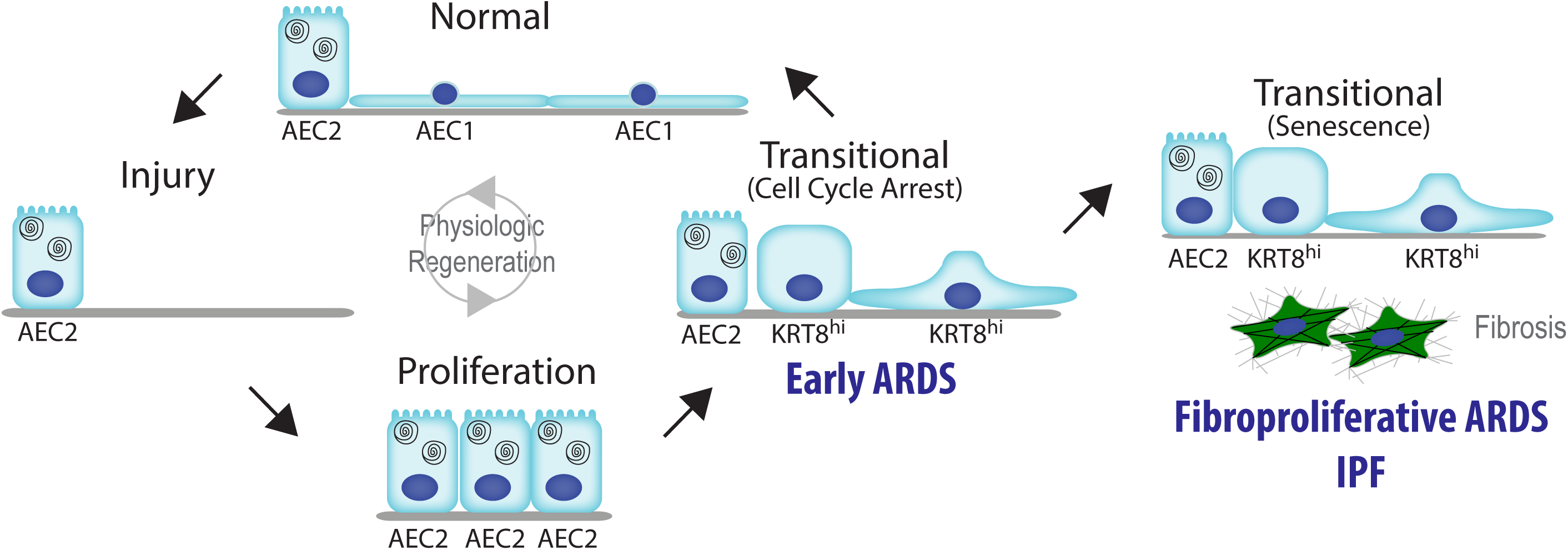

