## Supplemental Table 1 for "Fatal COVID-19 and non-COVID-19 Acute Respiratory Distress Syndrome is Associated with Incomplete Alveolar Type 1 Epithelial Cell Differentiation from the Transitional State Without Fibrosis"

Supplemental Table 1. Clinical Characteristics of COVID-19 ARDS Patients

| Patient | Age | Sex | BMI (kg/m^2^) | Initial Presentation | Comorbidities | Symptom Onset to Hospitalization (Days) | Symptom Onset to Intubation (Days) | Symptom Onset to Death (Days) | Radiologic Findings (Bilateral) | | Pharmacotherapies | Complications | | | Cause of Death |
| --- | --- | --- | --- | --- | --- | --- | --- | --- | --- | --- | --- | --- | --- | --- | --- |
|  |  |  |  |  |  |  |  |  | Ground Glass | Consolidation |  | Bacterial Pneumonia | Multiorgan failure | Other |  |
| 1 | 37 | M | 34.2 | Cough, fever, dyspnea | Type 2 diabetes mellitus, obesity | 1 | 5 | 10 | Yes | Yes | Hydroxychloroquine, prednisone, empiric antibiotics | No | Yes – renal, cardiac | -- | Multiorgan failure |
| 2 | 46 | M | 46.5 | Cough, fever, dyspnea, diarrhea | Type 2 diabetes mellitus, obesity | 3 | 9 | 16 | Yes | No | Hydroxychloroquine, empiric antibiotics | No | Yes – renal, hepatic | -- | Progressive hypoxic respiratory failure |
| 3 | 53 | M | 36.6 | Chest pain, dyspnea | Hyperlipidemia | 1 | 1 | 12 | Yes | No | Hydroxychloroquine, empiric antibiotics, sarilumab | No | Yes – renal, hepatic | Aortic dissection | Cardiac (PEA) arrest |

| Patient | Ventilator Settings | Pulmonary Mechanics | Arterial Blood Gas  (pH/pCO2/pO2/HCO3) | Rescue Modalities | | |
| --- | --- | --- | --- | --- | --- | --- |
|  |  |  |  | Prone Positioning | Paralytics | ECMO |
| 1 | Mode: VC/AC  PEEP: 26 cmH2O  FiO2:100%  Rate: 22  TV: 6 mL/kg IBW | Peak Inspiratory Pressure: 43 cmH2O  Plateau Pressure: 39cmH2O  Permissive Hypercapnia: no  P/F Ratio: 68 | 7.33 / 44/ 68 / 22.3 | Yes | Yes | Yes |
| 2 | Mode: VC/AC  PEEP: 24 cmH2O  FiO2: 100%  Rate: 36  TV: 5.3 mL/kg IBW | Peak Inspiratory Pressure: 46 cmH2O  Plateau Pressure: 43cmH2O  Permissive Hypercapnia: yes  P/F Ratio: 68 | 7.25 / 61 / 68 / 26 | No | Yes | No |
| 3 | Mode: PC/AC  PEEP: 22 cmH2O  IPAP: 41 cmH2O  FiO2: 100% | Tidal Volume: 8cc/kg IBW  Permissive Hypercapnia: no  P/F Ratio: 74 | 7.11 / 50 / 74 / 15.3 | Yes | Yes | No |
