## Supplemental Table 2 for "Fatal COVID-19 and non-COVID-19 Acute Respiratory Distress Syndrome is Associated with Incomplete Alveolar Type 1 Epithelial Cell Differentiation from the Transitional State Without Fibrosis"

Supplemental Table 2. Clinical Characteristics of non-COVID-19 ARDS Patients

| Patient | Age | Sex | BMI (kg/m^2^) | Initial Presentation | Comorbidities | Symptom Onset to Hospitalization (Days) | Symptom Onset to Intubation (Days) | Symptom Onset to Death (Days) | Radiologic Findings  (Bilateral) | | Pharmacotherapies | Complications | | | Cause of Death |
| --- | --- | --- | --- | --- | --- | --- | --- | --- | --- | --- | --- | --- | --- | --- | --- |
|  |  |  |  |  |  |  |  |  | Ground Glass | Consolidation |  | Bacterial Pneumonia | Multiorgan failure | Other |  |
| 4 | 40 | M | 21.1 | Abdominal distension, jaundice, shortness of breath | Alcoholic cirrhosis | 3 | N/A | 11 | Yes | No | Antibiotics, diuretics, steroids | No | Yes – renal | -- | ARDS |
| 5 | 37 | M | 34.4 | Chest pain | Hypertension | 0 | 0 | 14 | Yes | No | Antibiotics, vasopressors, anticoagulation, steroids | Yes | Yes – renal, cardiac, hepatic | Aortic dissection | ARDS and shock |
| 6 | 66 | M | 22.5 | Fever, shortness of breath | Former IV drug abuse, hepatitis C cirrhosis | 1 | 1 | 3 | Yes | No | Antibiotics, vasopressors, steroids | Yes | No | No | ARDS |

| Patient | Ventilator Settings | Pulmonary Mechanics | Arterial Blood Gas  (pH/pCO2/pO2/HCO3) | Rescue Modalities | | |
| --- | --- | --- | --- | --- | --- | --- |
|  |  |  |  | Prone Positioning | Paralytics | ECMO |
| 4 | Not applicable – 2L supplemental oxygen while on hospice care | -- | -- | No | No | No |
| 5 | Mode: PC-IMV/BiLevel  PEEP: 12 cmH2O  FiO2: 100%  Rate: 26  TV: 4.8 mL/kg IBW | Peak Inspiratory Pressure: 34 cmH2O  Plateau Pressure: 32 cmH2O  Permissive Hypercapnia: yes  P/F Ratio: 81 | 7.04 / 55 / 81 / 14.1 | No | Yes | Yes |
| 6 | Mode: PC-IMV/BiLevel  PEEP: 10 cmH2O  FiO2: 100%  Rate: 24  TV: 4.2 mL/kg IBW | Peak Inspiratory Pressure: 27 cmH2O  Plateau Pressure: 26 cmH2O  Permissive Hypercapnia: no  P/F Ratio: 58 | 7.24 / 44 / 58 / 18.4 | No | Yes | No |
