## Supplemental Figure Legends for "Fatal COVID-19 and non-COVID-19 Acute Respiratory Distress Syndrome is Associated with Incomplete Alveolar Type 1 Epithelial Cell Differentiation from the Transitional State Without Fibrosis"

**Supplemental Figure 1. Progression of Bilateral Infiltrates Consistent with Pulmonary Edema in a COVID-19 Patient.** Chest radiographs from a patient (patient #1) with COVID-19 ARDS taken on the indicated day of hospitalization. The patient died on hospital day 9.

**Supplemental Figure 2. Massive Loss of Mature AEC2s Observed in All COVID-19 ARDS Patients.** Immunostaining for the AEC2 marker proSPC revealed vast areas devoid of proSPC+ AEC2s with rare areas of proSPC+ AEC2s remaining. Scale bar = 100 µm. n=3.

**Supplemental Figure 3. AEC2 Hyperplasia is Rare in COVID-19 ARDS.** Immunostaining for the AEC2 marker proSPC revealed rare areas of hyperplastic mature AEC2s. Scale bar = 50 µm. n=3.

**Supplemental Figure 4. KRT8^hi^ Transitional Cells were Abundant throughout Lungs of All COVID-19 ARDS Patients.** Immunostaining for KRT8 revealed KRT8^hi^ transitional cells that were cuboidal and isolated (open yellow arrowheads), occurring in pairs (closed yellow arrowheads) or hyperplastic, suggesting recent cell division, partially spread (open white arrowheads), or, in rare cases, flat (closed white arrowheads). Occasional KRT8^hi^ transitional cells were hypertrophic and bizarrely shaped (closed orange arrowheads). Scale bar = 200 μm. n=3. Some images also shown in Fig. 2.

**Supplemental Figure 5. Normal AEC1s and AEC2s Express Much Lower Levels of KRT8 than Transitional Cells.** Lung sections were stained for KRT8. Expression of KRT8 in normal AEC1s (open arrowheads) and AEC2s (closed arrowheads) was observed with very high exposure times. Equal exposure times were used for both panels, which allowed low levels of K8 expression in normal lungs to be detected but resulted in oversaturation in transitional cells in COVID-19 lungs due to high levels of K8 expression. Scale bar = 50 μm. n=3.

**Supplemental Figure 6. KRT8^hi^ Transitional Cells do not Express the Club Cell Marker CCSP.** Club cells in normal airways express CCSP. Transitional cells in COVID-19 ARDS do not express CCSP. Scale bars = 50 µm. n=3.

**Supplemental Figure 7. Abundance of KRT8^hi^ Transitional Cells with Incomplete Differentiation into Mature AEC1s Observed in All COVID-19 ARDS Patients.** KRT8^hi^ transitional cells filled gaps denuded of AEC1s, were cuboidal, partially spread, or approaching a flat AEC1 morphology but typically did not express AEC1 markers. These findings were thought to represent ongoing organized, physiologic, albeit incomplete, AEC1 regeneration. Although all morphologies were present in all patients, Patient 3 tended to have more bizarrely-shaped cells. Scale bars = 100 µm. n=3. Some images also shown in Fig. 2.

**Supplemental Figure 8. Histology of All COVID-19 ARDS Patients is Acute DAD.** Lung sections were stained with hematoxylin and eosin. Patient 1: Early acute diffuse alveolar damage with intra-alveolar fibrin (star) and degenerating AECs (arrows) within alveolar space (100X). Patient 2: Acute diffuse alveolar damage with AEC hyperplasia (arrows) and hyaline membranes (arrowhead) (200X). Patient 3: Acute diffuse alveolar damage with AEC hyperplasia (arrows) (200X). Some images also shown in Fig. 1.

**Supplemental Figure 9. Transitional Cells and AEC1s Express Lower Levels of the SARS-CoV-2 Receptor ACE2.** Single cell RNA sequencing datasets from two mouse model of physiologic regeneration, LPS ^20^ and pneumonectomy (PNX) ^23^, and normal and IPF human lung ^31, 32^ were interrogated for expression of the SARS-CoV-2 receptor ACE2. Transitional cells and AEC1s express much lower levels of ACE2 than AEC2s.

**Supplemental Figure 10. Histology of All Non-COVID-19 ARDS Patients is Acute DAD.** Lung sections were stained with hematoxylin and eosin. Scale bar = 100 µm.

**Supplemental Figure 11. AEC1 Injury, AEC2 Proliferation, Abundant Transitional Cells, and Ongoing but Incomplete AEC1 Differentiation in Non-COVID-19 ARDS.** Lung sections were stained for HTI-56, KRT8, and proSPC. A) AEC1 loss. Scale bar = 50 µm. B) AEC2 hyperplasia. Scale bar = 50 µm. C) Abundance of transitional cells. Scale bar = 50 µm. D) KRT8^hi^ transitional cells fill gaps denuded of AEC1s on alveolar septa. Scale bar = 50 µm. Some are cuboidal (closed yellow arrowheads); some are partially spread, some approach a flat AEC1 morphology but do not express AEC1 markers (open white arrowheads), and rare cells express AEC1 markers (closed white arrowhead). Flat cells that express AEC1 markers but not KRT8 (open yellow arrowhead) were interpreted as native AEC1s that were not damaged during lung injury. Findings suggest ongoing organized, albeit incomplete, AEC1 differentiation. Scale bar = 50 µm. n=3.

**Supplemental Fig. 12. AEC2s and Transitional Cells in All Non-COVID-19 ARDS Patients.** Lung sections were stained for KRT8 and proSPC. All patients demonstrated abundant transitional cells and AEC2s. Occasional proSPC+ KRT8^hi^ cells were observed in non-COVID ARDS lungs (arrowheads). Scale bar = 100 µm. n=3.

**Supplemental Figure 13. Absence of Fibrosis in Early Non-COVID-19 ARDS.** A) Trichrome highlighted (in blue) basement membranes and vascular adventitia in normal and non-COVID-19 ARDS lungs and collagen deposition with marked fibrosis in IPF. Scale bar = 200 µm. B) Immunostaining demonstrated abundant myofibroblasts in IPF (asterisk) but not non-COVID-19 ARDS. Scale bars = 50 µm. n=3.

**Supplemental Figure 14. Transitional Cells in IPF but not Non-COVID-19 ARDS are Senescent.** Lung sections were immunostained. KRT8^hi^ transitional cells in IPF but not non-COVID-19 ARDS express the highly specific marker of senescence CDKN2A/p16. Scale bars = 50 µm. n=3.

**Supplemental Figure 15. Proposed Lineage Trajectories in Physiologic Alveolar Regeneration, Early ARDS, and Fibrosis.** During physiologic regeneration after lung injury resulting in AEC death, surviving AEC2s proliferate, then exit the cell cycle and transiently adopt the transitional state but ultimately differentiate into mature AEC1s. In early ARDS, after AEC death, surviving AEC2s proliferate, then exit the cell cycle and adopt the transitional state, and are in the process of organized AEC1 differentiation, but incomplete differentiation is associated with ongoing barrier permeability, pulmonary edema, ventilator dependence and mortality. In fibroproliferative ARDS and IPF, after AEC death, surviving AEC2s proliferate, then exit the cell cycle and adopt the transitional state, but evolve into a permanent state of cell cycle arrest, or senescence, and fibrosis ensues. Adapted with permission of the American Thoracic Society from Jiang P et al, Ineffectual Type 2-to-Type 1 Alveolar Epithelial Cell Differentiation in Idiopathic Pulmonary Fibrosis: Persistence of the KRT8hiTransitional State. Am J Respir Crit Care Med. 2020 Jun 1;201(11):1443-1447. Copyright © 2021 American Thoracic Society. All rights reserved. The American Journal of Respiratory and Critical Care Medicine is an official journal of the American Thoracic Society. Readers are encouraged to read the entire article for the correct context [here](https://www.ncbi.nlm.nih.gov/pmc/articles/PMC7258651/). The authors, editors, and The American Thoracic Society are not responsible for errors or omissions in adaptations.
